# Genetically-linked simultaneous overexpression of multiple herbicide-metabolizing genes for broad-spectrum resistance in an agricultural weed *Echinochloa phyllopogon*

**DOI:** 10.1101/2022.01.02.474499

**Authors:** Hiroe Suda, Tomomi Kubo, Yusuke Yoshimoto, Keisuke Tanaka, Satoru Tanaka, Akira Uchino, Satoshi Azuma, Makoto Hattori, Takuya Yamaguchi, Masahiro Miyashita, Tohru Tominaga, Satoshi Iwakami

**Affiliations:** Graduate School of Agriculture, Kyoto University, Kitashirakawa-Oiwake-cho, Sakyo-ku, Kyoto 606-8502, Japan; NODAI Genome Research Center, Tokyo University of Agriculture, 1-1-1 Sakuragaoka, Setagaya-ku, Tokyo 156-8502, Japan; Faculty of Regional Environment Science, Tokyo University of Agriculture, 1-1-1 Sakuragaoka, Setagaya-ku, Tokyo 156-8502, Japan; Central Region Agricultural Research Center, National Agriculture and Food Research Organization, Tsu 514-2392, Japan; Niigata Agricultural Research Institute Crop Research Center, 857 Nagakura-machi, Nagaoka, Niigata 940-0826, Japan; Biotechnology Research Center and Department of Biotechnology, Toyama Prefectural University, 5180 Kurokawa, Imizu, Toyama 939-0398, Japan

**Author notes:** Corresponding Author: Satoshi Iwakami.

**Keywords:** cytochrome P450, herbicide resistance, evolution, cross-resistance, NIH-shift

## Abstract

- Previous research unveiled that the overexpression of catalytically promiscuous CYP81A cytochrome P450s underlies the multiple-herbicide resistance (MHR) in a Californian population of *Echinochloa phyllopogon*. However, it does not fully accommodate the resistance to diverse herbicides in MHR *E. phyllopogon* although the genetic inheritance of MHR was suggested as under a single gene control.
- We investigated the high-level resistance to diclofop-methyl in MHR *E. phyllopogon*. Detailed diclofop-methyl metabolism was analyzed, followed by gene expression study and functional characterization of P450 genes. The generality of the MHR mechanism was investigated using another MHR line.
- The MHR line rapidly produced two distinct hydroxylated-diclofop-acid, only one of which was the major metabolite produced by CYP81A12/21. Gene expression study identified the genetically linked overexpression of a novel gene *CYP709C69* with *CYP81A12/21* in the MHR line. The gene conferred diclofop-methyl resistance in plants and produced another hydroxylated-diclofop-acid in yeast. The activity was observed in some CYP709C in plants. Unlike the broad substrate-specificity in CYP81As, CYP709C69 showed narrow substrate-specificity. The overexpression of the CYP81A and CYP709C69 was also observed in another MHR line.
- The present findings establish a novel concept that genetically-linked simultaneous overexpression of herbicide-metabolizing genes enhances and broadens the profile of metabolic resistance in weeds.

## Introduction

Heavy reliance on herbicide for weed control in agricultural fields have led to the evolution of herbicide resistance in weeds (Powles & Yu, 2010). The devastating impact on agriculture can be brought about by enhanced herbicide-metabolism since weeds with the mechanism often exhibit a large scope of resistance to chemically unrelated herbicides (Yu & Powles, 2014). Cytochrome P450 monooxygenases (P450s) have been assigned as a major component of metabolism-based resistance in weeds. Studies over the past 20 years have clearly shown that some P450s especially from crops carry metabolizing activity to some herbicides, leading to the protection plants from herbicide injuries (Dimaano & Iwakami, 2021). The involvement of P450s in weed resistance is beginning to be understood at the molecular level, but it has not been fully outlined. In particular, how multiple-herbicide resistance (MHR) is realized needs to be studied in more detail.

*Echinochloa phyllopogon* (syn. *Echinochloa oryzicola*) is an allotetraploid (2x=4x=36) and predominantly self-fertilizing noxious weed in rice cultivation (Yamasue, 2001). The first case of resistance in *E. phyllopogon* was reported from California in the late 90s’ where the resistant populations exhibited resistance not only to the herbicides used in the fields at that time but also to many other herbicides commercialized later (Nandula *et al*., 2019). Intensive analyses of this MHR population have led to the discovery that the overexpression of two CYP81A P450s is responsible for the cross-resistance to many herbicides such as acetolactate synthase (ALS) inhibitors (Iwakami *et al*., 2014a; Dimaano *et al*., 2020), acetyl-CoA carboxylase (ACCase) inhibitors (Iwakami *et al*., 2019), a 1-deoxy-d-xylulose 5-phosphate synthase (DXS) inhibitor (Guo *et al*., 2019), and a synthetic auxin (Chayapakdee, 2019). The two P450s, CYP81A12 and CYP81A21, exhibit broad substrate specificity, i.e. catalytic promiscuity, against diverse herbicide chemistries, accommodating the resistance to multiple herbicides in the *E. phyllopogon* population (Dimaano *et al*., 2020). Through genetic segregation study, the overexpression of the two P450s was suggested to be controlled by an unidentified single element (Iwakami *et al*., 2014a). Recently, similar cases have been reported in other weeds, *Lolium rigidum* and *Echinochloa crus-galli*, where multiple-herbicide resistance is also mediated by the overexpression of herbicide-metabolizing CYP81A P450s (Han *et al*., 2021; Pan *et al*., 2022). These studies provided molecular evidence that the overexpression of promiscuous enzyme at least partly accommodates MHR that is often accompanied in weeds with enhanced herbicide metabolism.

While the identification of the responsible genes advanced the understanding of metabolic resistance in weeds, MHR in the *E. phyllopogon* population is not fully understood, among which is the high-level resistance to ACCase inhibitor diclofop-methyl. While MHR line exhibit high-level resistance to diclofop-methyl, the diclofop-methyl metabolizing activity of the overexpressed P450 CYP81A12/21 was not high enough (Iwakami *et al*., 2019). Considering that diclofop-methyl resistance is inherited following the Mendelian segregation ratio as in other herbicides (Iwakami *et al*., 2019), we reasoned that an additional element specific to diclofop-methyl metabolism underlies the high-level diclofop-methyl resistance. Here, we report that the overexpression of a novel diclofop-methyl metabolizing P450, CYP709C69, is involved in high-level resistance to diclofop-methyl. In contrast to the catalytic promiscuity observed in CYP81As, CYP709C69 exhibited little metabolic activity to other herbicides except that it may be involved in the activation of a DXS inhibitor clomazone. This study highlights that genetically-linked simultaneous overexpression of herbicide-metabolizing P450s with different enzymatic properties is involved in MHR in weeds. The concept is distinct from the accumulations of multiple genes for broad or high-level resistance and serves as a model for future studies of metabolic resistance in weeds.

## Materials and Methods

### Origin of *E. phyllopogon*

Sensitive (S) and MHR lines (i.e. 401 and 511, respectively) originated from the Sacramento Valley in California (Fischer *et al*., 2000; Tsuji *et al*., 2003) were used. The F6 progeny of the two lines was derived from a single seed descent method (Iwakami *et al*., 2014a). The herbicide responses of F6 lines were evaluated previously (Iwakami *et al*., 2014a; Guo *et al*., 2019; Iwakami *et al*., 2019; Chayapakdee *et al*., 2020). Seeds of suspected S (Eoz1804) and resistant (Eoz1814) populations were collected at the paddy fields in Niigata prefecture in 2017. A plant of each population was cultivated and subjected to self-pollination, which was repeated twice before use. The full-length sequences of ALS genes (two copies) and carboxytransferase domain of ACCase genes (four copies) were amplified in a copy-specific way and directly sequenced as previously described (Iwakami *et al*., 2012).

### Analysis of diclofop metabolites in plants

*E. phyllopogon* were cultured hydroponically to around 2.5 leaf-stage at 25°C with a 12-h photoperiod (approximately 300 µmol m^-2^s^-1^) as described previously (Iwakami *et al*., 2019). The shoots of hydroponically cultured *E. phyllopogon* (2.5 leaf stage) were dipped in 30 µM diclofop-methyl solution supplemented with Tween 20 (0.01%) for 30 min. The shoots of 10 plants (approximately 300 mg) were collected at 0, 1, 3, and 6 h after application. On collection, they were rinsed in distilled water containing 20% methanol and 0.2% Triton X-100, and snap-frozen in liquid nitrogen. The plant tissue was ground into powder in liquid nitrogen with a mortar and pestle and extracted with 6 ml of 80% cold methanol, followed by centrifugation at 9,000 *g* for 10 min at 4°C. The pellet was further extracted twice with 2 ml of 80% cold methanol. The extract was combined, evaporated to dryness, and dissolved in 1 ml of 0.1 M sodium acetate buffer (pH 5.0). The solution was treated with 0.3 mg of almond β-glucosidase (BGH-101, Toyobo, Osaka, Japan) and incubated for 24 h at 37°C. After the addition of 1 ml of acetonitrile, the solution was vortexed, left for 15 min at 4°C, and then centrifuged at 1,700 *g* for 15 min. The supernatant was collected and evaporated to remove acetonitrile. The sample was loaded onto a Sep-Pak C18 cartridge (Waters, Tokyo, Japan) and washed with 10 ml of water containing 0.1% formic acid, followed by elution with 4 ml of acetonitrile. The fraction eluted with acetonitrile was evaporated to dryness and redissolved in acetonitrile (0.5 ml/300 mg of fresh weight of plant tissue), which was subjected to mass spectrometric analysis.

### Mass spectrometric analysis

Hydroxylated metabolites of diclofop-acid were analyzed using a liquid chromatography-tandem mass spectrometer (Shimadzu LCMS-8030, Kyoto, Japan) equipped with an electrospray ionization source. The MS parameters used were as follows: interface voltage of 4.5 kV, desolvation line temperature of 250°C, heat block temperature of 400°C, nebulizing gas (N_2_) of 2.0 l min^-1^, and drying gas at 15 l min^-1^. Multiple reaction monitoring (MRM) was used to quantitate the hydroxylated metabolites of diclofop-acid under the following conditions: negative ion mode with MRM transition of *m/z* 341 to *m/z* 269 (CE= 15 V, Q1 Pre Bias= 15 V, Q3 Pre Bias= 19 V). Separation of analytes was carried out using a reversed-phase column, TSK gel ODS-100V (2 mm ID x 150 mm, 3 µm, Tosoh, Tokyo, Japan). The column was eluted with a linear gradient from 30 to 90% mobile phase B (0.1% formic acid in acetonitrile) in mobile phase A (0.1% formic acid in water) for 20 min at a flow rate of 0.2 ml min^-1^ at 40°C. The injection volume was 5 μL.

### RNA extraction and cDNA synthesis

Total RNA was extracted using an RNeasy Plant Mini Kit (Qiagen, Tokyo, Japan) followed by the application of TURBO DNA-*free* kit (Thermo Fisher Scientific, Tokyo, Japan) to eliminate genomic DNA. Total RNA (1 μg) was reverse-transcribed using ReverTra Ace (Toyobo, Osaka, Japan) according to the manufacturer’s instructions.

### RNA-seq analysis

RNAs extracted from the shoot of 2.5-leaf stage plants of S and MHR lines were used for RNA-seq analysis with four biological replications. RNA-Seq libraries were prepared with TruSeq RNA Library Preparation Kit v2 (Illumina, USA), followed by pair-end sequencing on HiSeq 2500 sequencer (Illumina). Reads were filtered using TRIMMOMATIC (ver 0.36) (Bolger *et al*., 2014) with the following options: TRAILING:25, SLIDINGWINDOW:4, MINLEN:80. The reads were mapped against coding sequences of the draft genome of *E. phyllopogon* (Ye *et al*., 2020) using bowtie2 (ver 2.3.5.1) with the following options: ‘-X 900–very-sensitive–no-discordant–no-mixed–dpad 0–gbar 99999999’. The mapped reads were counted using RSEM (Li & Dewey, 2011). Differential expression analysis was performed using edgeR (ver 3.28.0) with glmQLFTest (Robinson *et al*., 2010) in R (R Core Team, 2016). Contigs were annotated by BLASTX (Blast+ v.2.9.0) (Camacho *et al*., 2009) against rice protein sequences (*Oryza sativa* MSU release 7) and rice P450 protein sequences in Cytochrome P450 Homepage (https://drnelson.uthsc.edu/).

### Real-time PCR

Real-time PCR was performed as described previously (Tanigaki *et al*., 2021) using primers listed in Table S1. Some of the primer sets were developed elsewhere (Czechowski *et al*., 2005; Iwakami *et al*., 2014b; Tanigaki *et al*., 2021). Eukaryotic translation initiation factor 4B (*EIF4B*) was used as an internal control gene and data was analyzed by the ΔΔCt method (Schmittgen & Livak, 2008).

### Isolation of P450 genes

The full-length sequence of each P450 gene was amplified from the cDNA of *E. phyllopogon* (line 511) using primers listed in Table S1 with KOD FX Neo (TOYOBO) or PrimeSTAR GXL DNA Polymerase (Takara). The amplicons were subcloned using pGEM-T Easy Vector Systems Kit (Promega, Tokyo, Japan) or Zero Blunt PCR Cloning Kit (Thermo Fisher Scientific, Tokyo, Japan) according to the manufacturer’s instructions.

### Plant transformation

The coding region of each P450 gene was cloned into pCAMBIA1390 vector with the In-Fusion DH Cloning Kit (TaKaRa, Kusatsu, Japan) or SLiCE reaction (Motohashi, 2015) using primers listed in Table S1. The binary construct was introduced into *Agrobacterium tumefaciens* strain EHA105 using the freeze and thaw method (Hofgen & Willmitzer, 1988). Rice calli (*Oryza sativa* cv. Nipponbare) were transformed as previously described (Toki, 1997; Iwakami *et al*., 2019). *Arabidopsis thaliana* (ecotype Columbia-0) was transformed by floral dip method (Clough & Bent, 1998) with the selection of hygromycin B (20 mg l^-1^). T3 homozygous lines were selected based on the segregation ratio.

### Herbicide sensitivity assay

Independently transformed calli were placed on N6D solid media supplemented with a herbicide as described previously (Iwakami *et al*., 2019). The calli were cultured at 30°C for 3 weeks. For the *A. thaliana* sensitivity assay, the surface-sterilized seeds were placed on Murashige and Skoog solid media (Murashige & Skoog, 1962). After three days of stratification at 4°C, the media were placed at a climate chamber with 22°C, 12-h photoperiod, and 70 μE m^-2^ s^-1^ light intensity for 12 days. In the case of the diclofop-methyl assay, the plates were kept in the chamber for 14 days. Chlorophyll was extracted and measured as described previously (Endo *et al*., 2021). The herbicides and the doses were summarized in Table S2.

The diclofop-methyl response of *E. phyllopogon* was determined as described previously (Iwakami *et al*., 2019). Briefly, hydroponically cultured plants were dipped in diclofop-methyl solution for 30 min as described above. The plants were further cultivated for nine days. Dose-response curves were drawn with the drc package (ver 3.0) (Ritz *et al*., 2015).

### Heterologous expression of P450s in yeast and whole-cell herbicide metabolism assay

Expression in yeast (*Saccharomyces cerevisiae*) was conducted using WAT11 strain and pYeDP60 vector system (Pompon *et al*., 1996). The coding regions of other P450s were cloned into the pYeDP60 vector with the yeast Kozak sequence inserted as described before (Iwakami *et al*., 2019). WAT11 was transformed using the lithium acetate method (Ito *et al*., 1983). A whole-cell assay for diclofop-methyl metabolism was performed according to Iwakami *et al*. (2019) with the modification of diclofop-methyl concentration to 300 μM. After incubation for 24 h, yeast cells were centrifuged at 13,500 *g* for 10 min. The supernatant was cleaned up using a Sep-Pak C18 cartridge as described above and subjected to liquid chromatography-tandem mass spectrometry (LC-MS/MS) analysis.

### Structure determination of diclofop metabolites

The yeast expressing *CYP81A12* or *CYP709C69* in whole-cell assay buffer (350 ml) was incubated with diclofop-methyl (300 μM) for 24 h. The culture was centrifuged at 1,500 *g* for 5 min to collect the supernatant. The yeast cells were resuspended in 1/10 volume of the buffer without diclofop-methyl. After incubation for an additional 12 h, the supernatant was collected by centrifugation, which was repeated twice. The whole procedure was repeated 30 times. After combining all the supernatants, the pH of the solution was adjusted to 1 by hydrochloric acid. The solution was extracted four times with diethyl ether (0.7 volume). After evaporating diethyl ether *in vacuo*, the resultant residue was dissolved in acetonitrile. High-performance liquid chromatography (HPLC) (LC-10ADvp and SPD-M10Avp, Shimadzu) separation was carried out on a reversed-phase column (TSK gel ODS-100, 4.6 mm ID x 250 mm, 5 µm, Tosoh, Tokyo, Japan) with the following conditions: linear gradient from 30 to 72% mobile phase B (0.1% formic acid in acetonitrile) in mobile phase A (0.1% formic acid in water) for 35 min at a flow rate of 0.8 ml/min at 40°C. The elution was monitored by UV absorbance at 215 and 280 nm. The fractions containing each hydroxylated metabolite of diclofop-acid were evaporated to dryness and dissolved in deuterated chloroform. ^1^H-nuclear magnetic resonance (NMR) spectra of each metabolite were recorded on a Bruker Ascend 400 MHz spectrometer (Billerica, MA, USA) with tetramethylsilane as an internal standard.

### Phylogenetic analysis of *CYP81A* and *CYP709C*

Nucleotide sequences of CYP709C subfamilies of rice were obtained from *Oryza sativa* MSU release 7 (Ouyang *et al*., 2007), and RAP-DB (Sakai *et al*., 2013). Deduced protein sequences were aligned with MAFFT (ver 7.475) (Katoh & Standley, 2013). The alignments were used to estimate evolutionary topology in MEGA X (Kumar *et al*., 2018) using Neighbor-Joining method. JTT matrix-based method was used to compute the evolutionary distances. Bootstrap values (1,000 replicates) are shown next to the branches.

### Synteny analysis

The genome of *E. haploclada* (Ye *et al*., 2020) was used for the synteny analysis. The local synteny around the *CYP81A*s and *CYP709C*s was visualized with clinker (ver 0.0.21) (Gilchrist & Chooi, 2021).

## Results

### Two distinct hydroxylated metabolites of diclofop-acid are highly accumulated in MHR *E. phyllopogon*

To gain clues for the high-level resistance to diclofop-methyl in the MHR line, we investigated the metabolism of diclofop-methyl in *E. phyllopogon*. Diclofop-methyl is a pro-herbicide that is activated by endogenous esterase as is the case for most aryloxyphenoxy-propanoate herbicides (Wenger *et al*., 2012). Following the de-esterification, the resultant active form diclofop-acid is inactivated by P450-mediated hydroxylation followed by further glucosylation (Fig. 1a) (Shimabukuro *et al*., 1979, 1987; McFadden *et al*., 1989). In the metabolism scheme, diclofop-acid hydroxylation is the rate-limiting step for diclofop-methyl inactivation in wheat, which is also suggested in other plants (Owen, 2000). Our previous study showed that both the MHR and S lines converted the majority of diclofop-methyl into diclofop-acid by 3 h after treatment for 30 min (Iwakami *et al*., 2019). In this study, we further analyzed the dynamics of putative OH metabolites of diclofop-acid in the extract of plants prepared as before (Fig. 1b). Two peaks with retention times of 11.3 and 12.4 min (M1 and M2, respectively), which correspond to putative OH metabolites, were detected by LC-MS/MS analysis using a MRM mode (Fig. 1c). The signal intensity of M2 was higher than that of M1 under this condition. In line with our study, a variety of hydroxylated diclofop-acid metabolites were reported in wheat (Tanaka *et al*., 1990; Zimmerlin & Durst, 1992), suggesting the presence of at least partially overlapping metabolic pathway for diclofop-methyl between wheat and *E. phyllopogon*.

**Figure 1.**
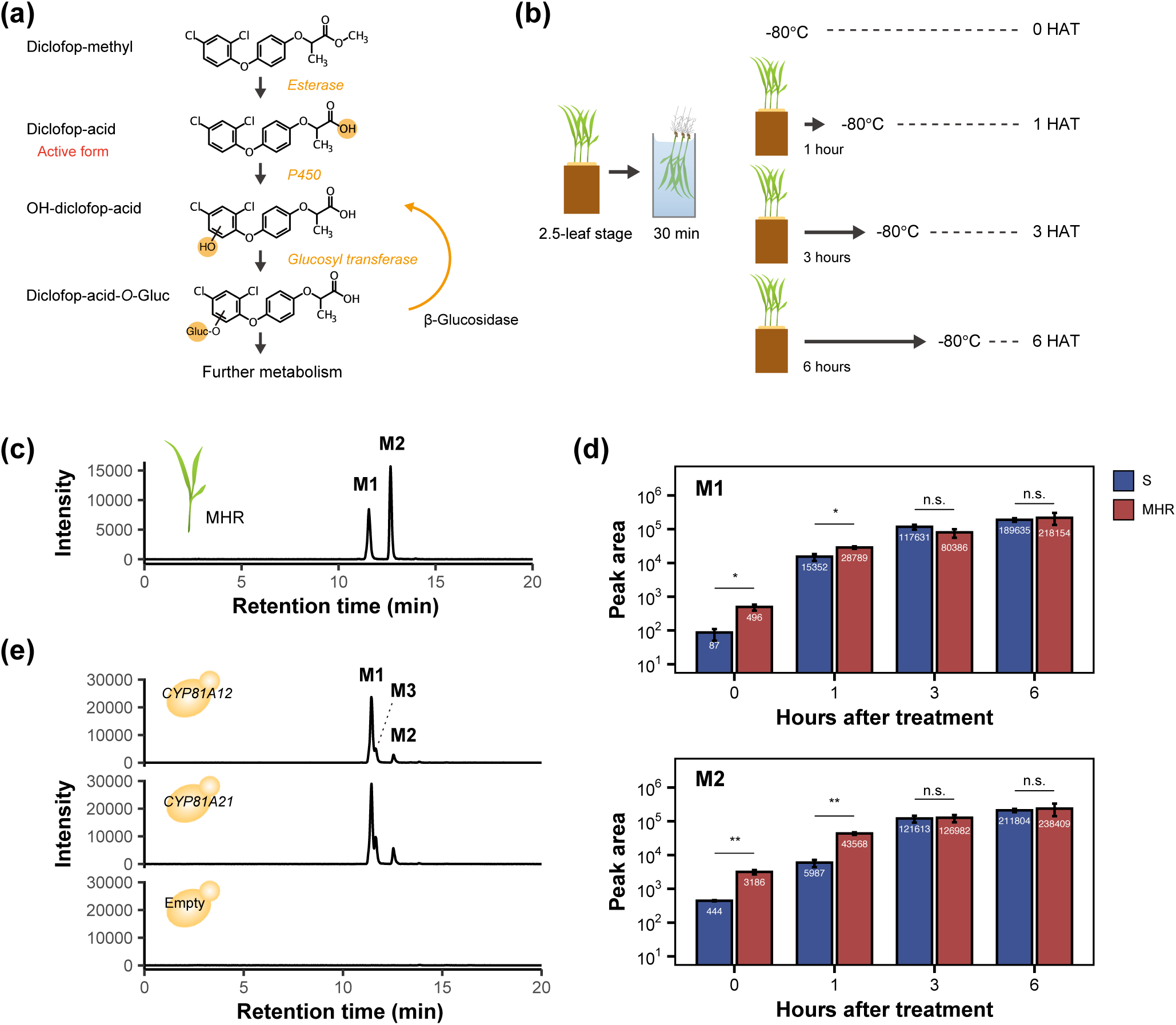
Characterization of diclofop-acid metabolism in multiple-herbicide resistant (MHR) *Echinochloa phyllopogon*. (a) Diclofop-methyl metabolism in plants. The enzymes estimated to be involved in each metabolism steps are shown. (b) Schematic representation of sample preparation for diclofop-methyl metabolism study. The shoots were dipped in diclofop-methyl solution (30 μM) for 30 min, which were collected and stored at 0, 1, 3, or 6 h after treatment (HAT). (c) Detection of OH-diclofop-acid in the MHR line of *E. phyllopogon* on LC-MS/MS. The extract of 3 HAT was analyzed. (d) Comparison of M1 and M2 metabolite amounts between sensitive (S) and MHR lines. Diclofop-acid-*O*-Gluc was converted to OH-diclofop-acid with β-glucosidase as shown in (a). The error bars, SEM. * and ** show significant differences of *P*< 0.05 and *P*< 0.01, respectively (Student’s *t*-test). (e) Whole cell diclofop metabolism assay using yeast harboring empty vector (pYeDP60), *CYP81A12,* or *CYP81A21*.

We next compared the amount of each hydroxylated metabolite between the MHR and S lines. To accurately evaluate the hydroxylation activity, possible glucosylated metabolites were converted into hydroxylated diclofop-acid by adding β-glucosidase to the extract of the diclofop-methyl treated plants (Fig. 1a), which was then subjected to comparative quantification between the lines. As a result, the amounts of both hydroxylated metabolites were higher in MHR plants at 0 and 1 HAT although no significant difference was observed at 3 and 6 HAT (Fig. 1d). Diclfop-methyl follows complexed metabolic pathways after glucosylation in plants (Shimabukuro *et al*., 1987; Han *et al*., 2013), which should have masked the difference of these metabolites between the lines later than 3 HAT.

To investigate whether the metabolites are produced by CYP81A12 and CYP81A21, the P450s responsible for the resistance to the majority of herbicides in the MHR line (Dimaano *et al*., 2020), we analyzed the whole cell diclofop-methyl metabolism of yeast carrying *CYP81A12/21*. In the yeast system, another peak (M3), which was not clearly separated from M1, was observed in addition to M1 and M2. Notably, the chromatographic profile of M1 and M2 differed between *E. phyllopogon* and yeast: the intensity of M2 was higher in *E. phyllopogon*, while that of M1 was prominently higher in yeast (Fig. 1e). The result suggests that an additional enzyme that catalyzes hydroxylation to form M2 is involved in diclofop-methyl resistance in the MHR line.

### Candidate genes for high-level diclofop-methyl resistance were identified by gene expression study

To identify the additional diclofop-hydroxylating enzyme, we performed RNA-seq analysis using the recently released draft genome of *E. phyllopogon* with 66,521 gene models (Ye *et al*., 2020). Among the gene models, we identified 549 putative P450 genes. A transcriptome comparison detected some P450 genes highly expressed in the MHR line. A closer look at the sequences of these P450 genes found that at least two of them (Contig817_pilon.380 and Contig 222_pilon.21) were misannotated due to the existence of similar genes next to each other (Fig. S1). Thus, we replaced the respective coding sequences with two correct sequences and mapped the reads to the revised transcriptome.

Finally, we detected 463 genes that were highly expressed in the MHR line (Fig. 2a, b, Fig. S2). Among the genes, 18 P450 genes were detected including previously reported multiple-herbicide metabolizing *CYP81A12* and *CYP81A21* taking the top four and five positions, respectively, among all the P450 genes in the MHR line (Fig. 2a, c). Four of them respectively encode a truncated protein: Contig168_pilon.233 (*CYP81A25P*), Contig1257_pilon.115 (*CYP81A19P*), Contig222_pilon21.b (*CYP704A*), and Contig774_pilon.23 (*CYP71E*); thus, they were excluded from further analyses.

**Figure 2.**
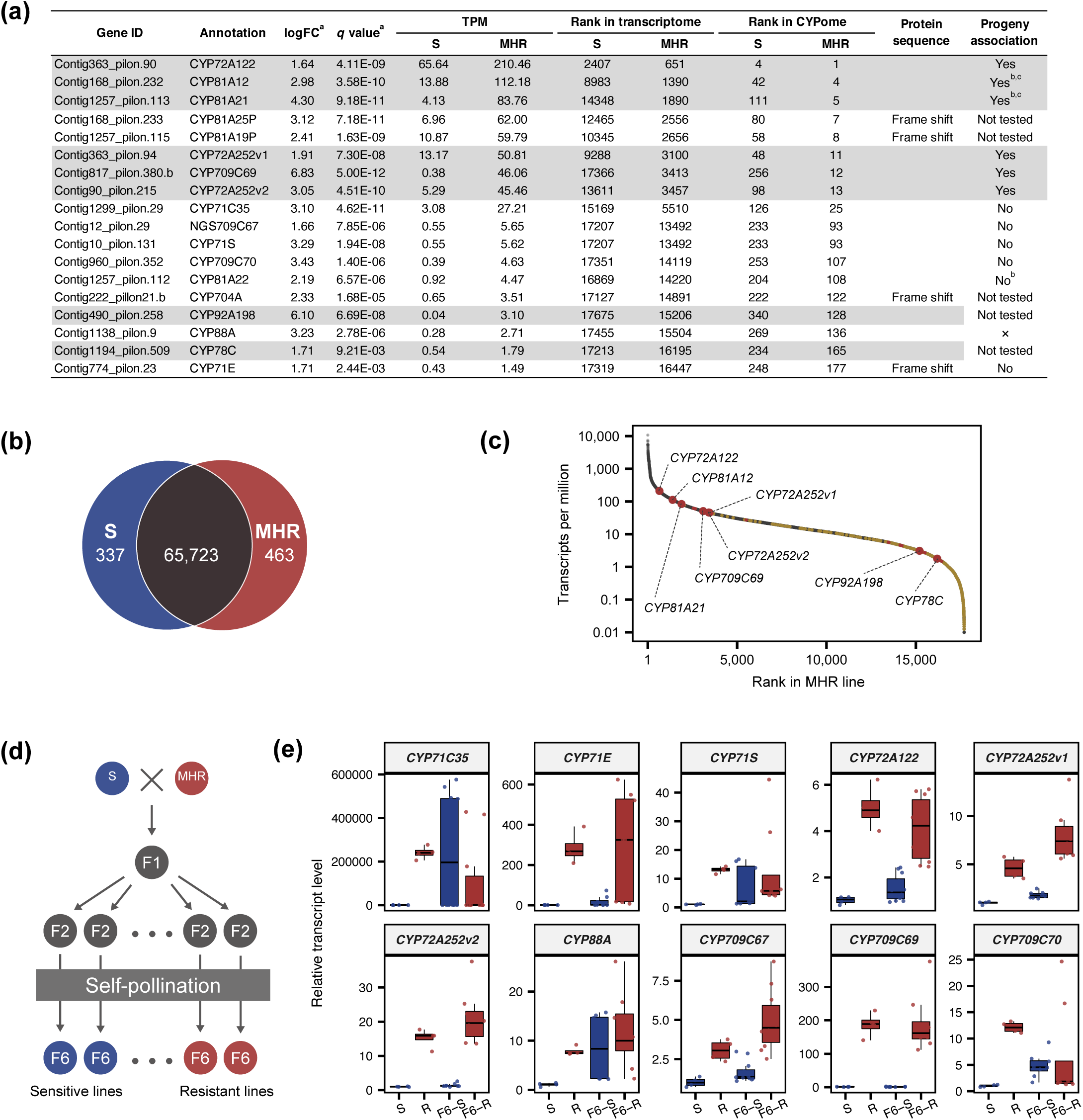
Candidate genes for high-level diclofop-methyl resistance in *Echinochloa phyllopogon* were identified by gene expression study. (a) The annotations of P450 genes that were upregulated in the MHR line. Genes are ordered according to MHR expression rank. Progeny association study was not conducted on Contig490_pilon.258 and Contig1194_pilon.509 since the appropriate primers were not designed. Highlight in gray indicates that the gene passed the tests. ^a^, calculated in edgeR. ^b^, Iwakami *et al*. (2014a). ^c^, Iwakami *et al*. (2019). (b) The number of differentially expressed genes. (c) Expression rank of P450 genes in the transcriptome of MHR line. P450s genes are shown in yellow and red. Red plots represent differentially expressed P450 genes. Larger Red plots are the genes shown in grey in Fig. 2a. (d) F6 lines used in the association study. (e) Progeny association of the mRNA levels in the shoots (four plants in bulk for each line) of the candidate P450 genes. mRNA levels were quantified by real-time PCR in 16 F6 lines that were previously characterized (Iwakami *et al*., 2019; Chayapakdee *et al*., 2020). The values for *CYP71C35* in the S and some F6 lines line were set as 40 cycles since the signal was not detected. F6-S, sensitive F6 lines; F6-R, resistant F6 lines.

The candidate overexpressed genes were further screened by real-time PCR for their co-segregation with MHR using the F6 recombinant inbred lines from a cross between the MHR and S lines (Fig. 2d). The analysis revealed that *CYP72A122*, *CYP72A252v1*, *CYP72A252v2*, and *CYP709C69* co-segregated with diclofop-methyl resistance (Fig. 2e), suggesting that they are the prime candidates for metabolite M2-producing P450(s). Thus, we subjected the four genes to subsequent functional characterization. We also tested the function of *CYP92A198* and *CYP78C* as we failed to evaluate them by real-time PCR due to failures in designing appropriate primers.

### Identification of novel diclofop-methyl metabolizing P450

For the evaluation of the diclofop-metabolizing activity of each gene, we used the rice transformation system as employed previously (Iwakami *et al*., 2019). We expressed the six candidate genes in rice calli with the pCAMBIA1390 vector, where the transgene was placed under the control of Cauliflower mosaic virus 35S promoter. The calli were tested for growth on the media supplemented with diclofop-methyl. Among the six genes, only *CYP709C69* conferred marked resistance at the lethal dose of diclofop-methyl (0.4 μM) to rice calli (Fig. S3a). No difference was observed in the coding sequence of *CYP709C69* between the S and MHR lines. When 12 independent transformed lines were subjected to diclofop-methyl, all the lines with *CYP709C69* stopped growing at 6 μM (Fig. 3a). The diclofop-methyl resistance level was equivalent to the cases of *CYP81A12* and *CYP81A21*. Decrease in diclofop-methyl sensitivity was also observed in a *A. thaliana* line transformed with *CYP709C69* as well as those with *CYP81A12* and *CYP81A21* (Fig. S4).

**Figure 3.**
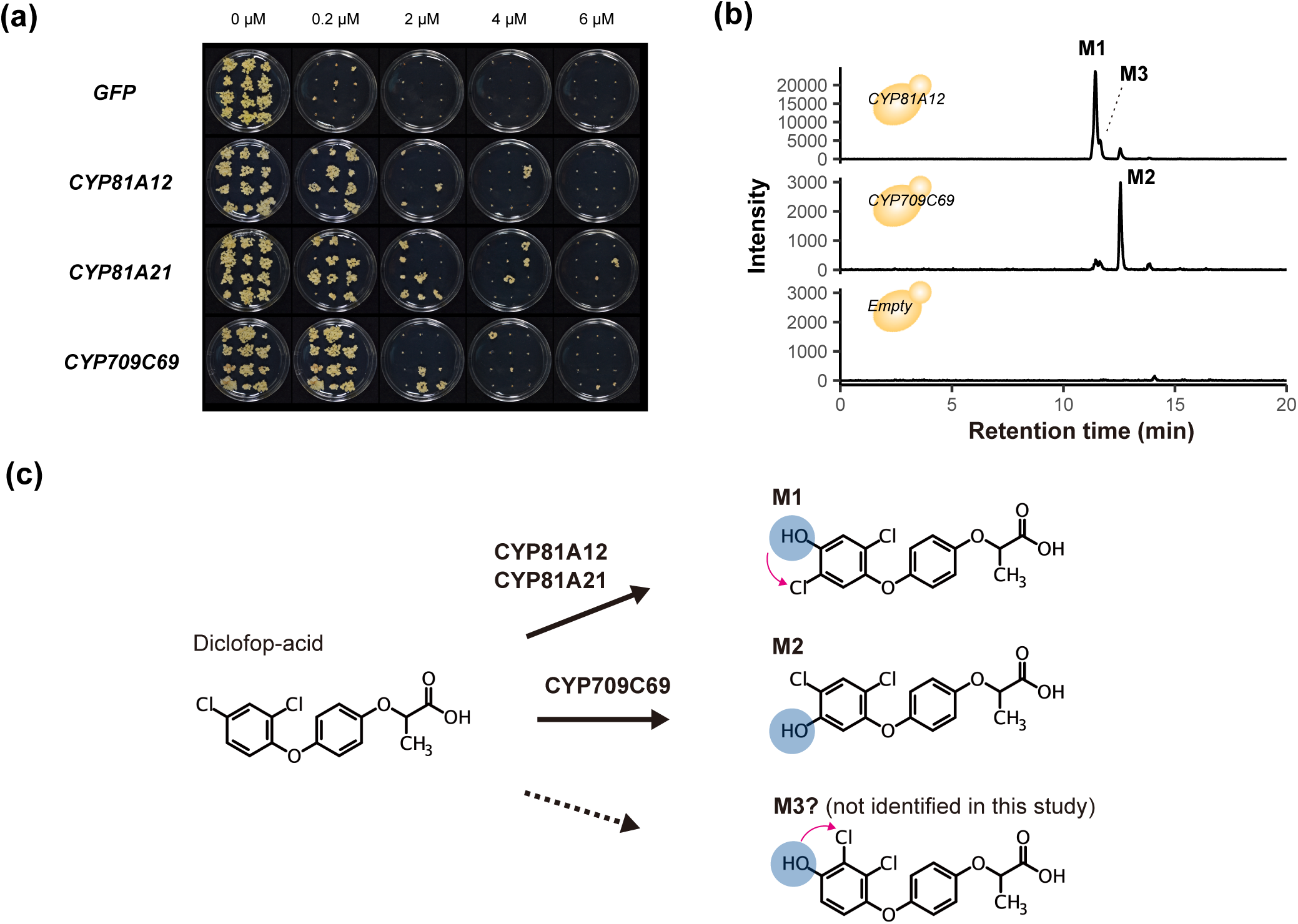
Identification of novel diclofop-methyl metabolizing P450. (a) Diclofop-methyl sensitivity of rice calli transformed with P450 genes of *Echinochloa phyllopogon.* Twelve independent calli were grown for three weeks. (b) Whole cell diclofop-methyl metabolism assay using yeast harboring empty vector (pYeDP60), *CYP81A12,* or *CYP709C69*. (c) The role of P450s in diclofop-acid hydroxylation. The structures of M1 and M2 were determined by nuclear magnetic resonance analysis. The arrows indicate migration of chlorine. The structure of M3 metabolite was estimated to have a 2,3-dichloro-4-hydroxyphenoxy moiety as reported in wheat (Tanaka *et al*., 1990).

We then transformed *CYP709C69* in yeast and investigated whole cell diclofop-methyl metabolism as performed previously (Iwakami *et al*., 2019). LC-MS/MS analysis revealed that M2 was produced as the main peak (Fig. 3b). To determine the hydroxylation position in the OH-metabolites of diclofop-acid, we purified M1 and M2 from yeast culture applied with diclofop-methyl using HPLC, which were then subjected to NMR analysis. Since the NMR structural analysis of three OH-metabolites of diclofop-methyl has been reported previously (Tanaka *et al*., 1990), we compared the chemical shift patterns in the aromatic regions of M1 and M2 with the reported data. The results showed that M1 has a similar pattern with the metabolite containing a 2,5-dichloro-4-hydroxyphenoxy moiety and M2 with that containing a 2,4-dichloro-5-hydroxyphenoxy moiety (Fig. S5). The position of one of the chlorine atoms in M1 is different from the original position, which is thought to be formed by intramolecular migration during enzymatic hydroxylation of the aromatic ring (so-called NIH-shift) as previously reported (Fig. 3c) (Tanaka *et al*., 1990). This implies that CYP81A12 and CYP81A21 mediate the NIH-shift reaction in diclofop metabolism. In this study, we could not determine the structure of M3 due to the limited amount available from CYP81A expressing yeast (Fig. 1e, 3b). Considering that wheat produces the hydroxylated diclofop with a 2,3-dichloro-4-hydroxyphenoxy moiety (Tanaka *et al*., 1990) in addition to the metabolites shown above, M3 may carry this moiety (Fig. 3c) although further experiments are required.

### Diclofop-acid metabolizing activity is partially conserved among CYP709C P450s

CYP709C was previously characterized in wheat, where the isolated CYP709C1 harbors sub-terminal hydroxylation activity of C18 fatty acids (Kandel *et al*., 2005) (Fig. 4a). Importantly, the *in vitro* microsome assay purified from yeast showed that CYP709C1 is inactive to diclofop-acid, raising a possibility that the observed diclofop-acid metabolizing activity in CYP709C69 may be specific to this P450. To investigate the functional conservation, we further tested other CYP709Cs in *E. phyllopogon* although no association with resistance was observed in the gene expression analyses (Fig. 2e). Since *E. phyllopogon* is an alletetraploid, a majority of genes have a close relative, as observed for CYP709C68 and Contig960_pilon.355, CYP709C70 and CYP709C71, CYP709C67 and Contig1380_pilon.11. However, the counterpart for CYP709C69 was missing in the genome (Fig. 4a). We first tested CYP709C68, one of the highly similar CYP709Cs to CYP709C69. The yeast expressing *CYP709C68* produced a small amount of M2 metabolite (Fig. 4b). The rice calli expressing *CYP709C68* exhibited no prominent difference in diclofop-methyl sensitivity (Fig. S3b), which may indicate its low activity against diclofop-methyl. Further study is required to determine its relative activity to CYP709C69 since the expression levels were not determined both in the yeast and rice calli. Four more genes, i.e., *CYP709C67*, *CYP709C68*, *CYP709C70*, *CYP709C71*, and Contig960_pilon.355 were also tested in rice calli system. However, the growth of the calli on diclofop-methyl media did not markedly differ from *GFP* expressing calli (Fig. S3b). The genome holds six more putative *CYP709C* genes (Contig1380_pilon.11, Contig12_pilon.1, Contig291_pilon.387, Contig1182_pilon.1, Contig291_pilon.396, and Contig120_pilon.313). However, the annotated protein sequences other than Contig1380_pilon.11 and Contig291_pilon.387 were too short (<331) for a canonical P450 sequence.

**Figure 4.**
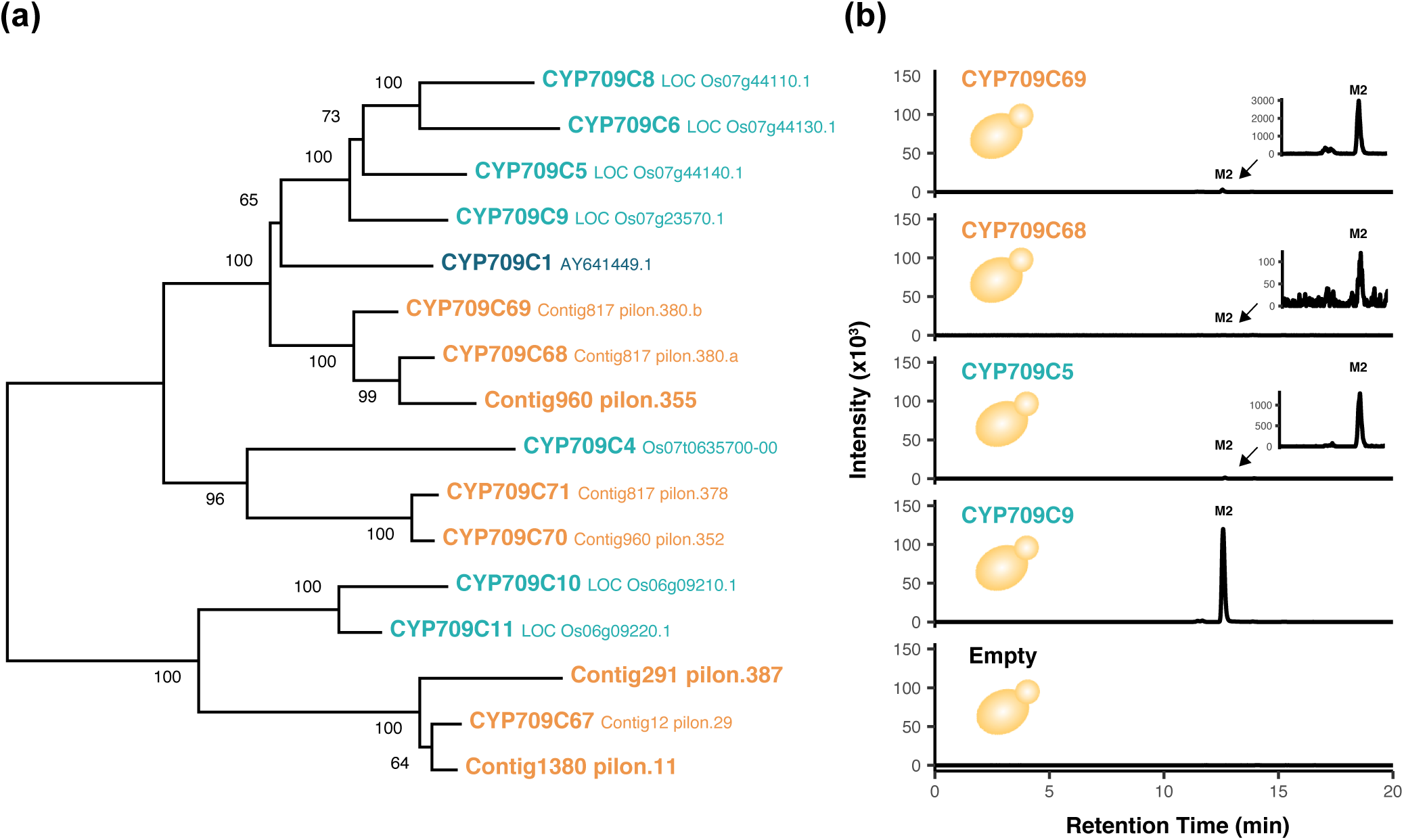
Diclofop-methyl metabolizing activity of CYP709Cs. (a) Phylogenetic tree of CYP709Cs in *Echinochloa phyllopogon* and rice, and CYP709C1 from wheat. CYP709C1 did not metabolize diclofop-acid in in vitro assay (Kandel *et al*., 2005). The gene ID or accession number is shown next to CYP name when CYP names are available. The annotation of Contig817_pilon.380.a and Contig817_pilon.380.b is shown in Fig. S1. (b) Whole cell diclofop-methyl metabolism assay using yeast harboring empty vector (pYeDP60), *CYP709C5, CYP709C9, CYP709C68,* and *CYP709C69*.

We further evaluated the function of the *CYP709C* gene in rice. Seven genes were annotated as *CYP709C* in rice (Fig. 4a). Of these, CYP709C5 and CYP709C9, which are relatively similar to CYP709C69, were tested in the yeast expression system. Both enzymes produced significant amount of M2 metabolite (Fig. 4b) although the activity cannot be directly compared with CYP709C69. Taken together, these results suggest that the ability to metabolize diclofop is at least partially conserved in CYP709C members.

### CYP709C69 has a narrow substrate specificity

To investigate the role of CYP709C69 in the metabolism of other herbicides, we tested the herbicide sensitivity of *CYP709C69*-expressing *A. thaliana*. We also used *CYP709C69*-expressing rice calli for the evaluation of its metabolizing activity to ACCase inhibitors. Among a total of 46 herbicides from at least 15 modes of action, a DXS inhibitor clomazone was the only herbicide where a marked alteration in the sensitivity of the transgenic plants was observed (Fig. S6). The *A. thaliana* lines expressing *CYP709C69* became more vulnerable to clomazone in inverse proportion to the expression level (Fig. 5a, b), in contrast to the *A. thaliana* transformed with *CYP81A*s (Guo *et al*., 2019). Clomazone is a pro-herbicide where P450 was proposed as the enzyme to convert 5-OH clomazone, and to further convert to the active form 5-keto clomazone (Fig. 5c) (Nandula *et al*., 2019). Thus, CYP709C69 may be involved in either of the sequential bio-activation reaction.

**Figure 5.**
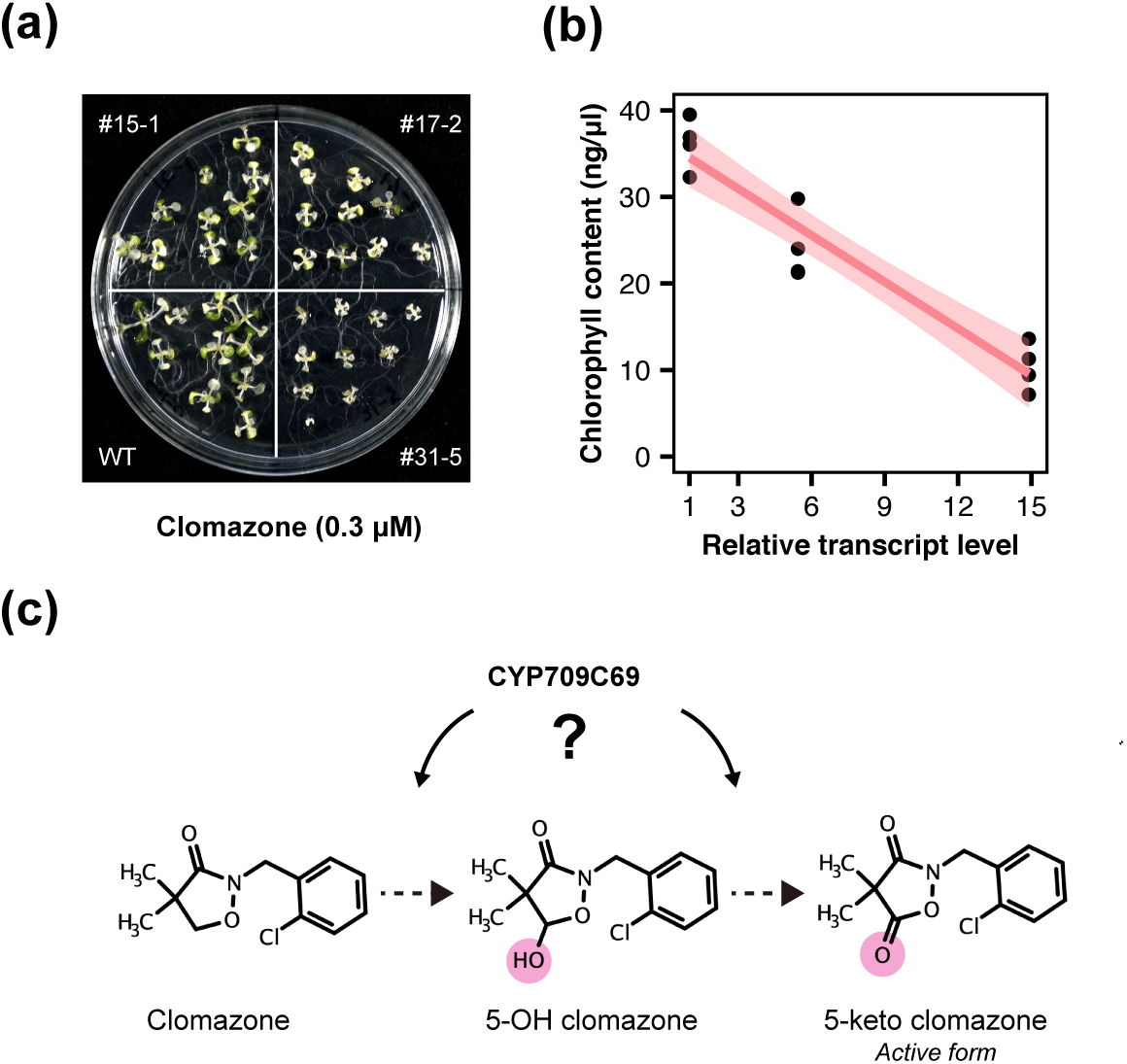
*CYP709C69* increased clomazone sensitivity in *Arabidopsis thaliana*. (a) Clomazone (0.3 μM) response of *A. thaliana* lines transformed with *CYP709C69*. The mRNA level of each line was evaluated in Fig. S4. (b) Association of transcript level of *CYP709C69* (n=4) and chlorophyll content in the 0.3 μM clomazone treated *A. thaliana* lines. Linear regression line is shown in pin with shaded regions representing the 95% confidence interval. (c) The pro-herbicide clomazone is converted to the active form 5-keto clomazone through 5-OH clomazone in plants. The involvement of P450s were suggested (Nandula *et al*., 2019). CYP709C69 may be involved in either of the reaction.

### Distantly located herbicide-metabolizing genes are simultaneously overexpressed

That the overexpression of the three P450 genes co-segregated in the F6 progenies suggest that a single causal element underlies the regulation. To further examine the possibility, we analyzed an additional 16 F6 lines, where the linkage of overexpression of *CYP81A12/21* with MHR was reported (Chayapakdee *et al*., 2020). In these additional lines, the overexpression of *CYP709C69* also linked with MHR (Fig. S7), supporting the simultaneous regulation of the three genes in the MHR line.

To gain insights into the mechanism of overexpression of the three herbicide-metabolizing P450 genes (*CYP81A12*, *CYP81A21*, and *CYP709C69*), we estimated the genomic loci of the three genes. Since the draft genome of *E. phyllopogon* is contig-based, the physical relationships of the three P450 genes in the chromosome are unclear. Thus, we allocated the loci of the three P450 genes in the chromosome-scale genome of diploid *Echinochloa* sp. (*E. haploclada*) (Ye *et al*., 2020). The *CYP81A* and *CYP709C* loci respectively correspond to chromosomes 1 and 3 in the *E. haploclada* genome (Fig. 6a). The synteny around the *CYP81A* regions (Contig168 and Contig1257) and *CYP709C* regions (Contig817 and Contig960) is highly conserved in the *E. haploclada* genome (Fig. 6b), implying that the contigs are fragments of homeologous chromosomes. Importantly, the synteny structure shows that *CYP81A12* and *CYP81A21* represent homeologous relationship as hinted previously from their phylogenetic relationships and herbicide-metabolizing functions (Iwakami *et al*., 2014a; Dimaano *et al*., 2020). A homeologous gene for *CYP709C69* was not identified on Contig960, which is consistent with the phylogenetic analysis (Fig. 4a). Considering that the MHR including diclofop-methyl resistance is inherited as a Mendelian trait (Iwakami *et al*., 2014a, 2019) and that the overexpression of the three genes is genetically linked (Fig. 2, Fig. S7), a single genetic element is suggested to be behind the MHR of the Californian population of *E. phyllopogon*. The causal element may directly or indirectly regulate the expression of the P450s (Fig. 6c).

**Figure 6.**
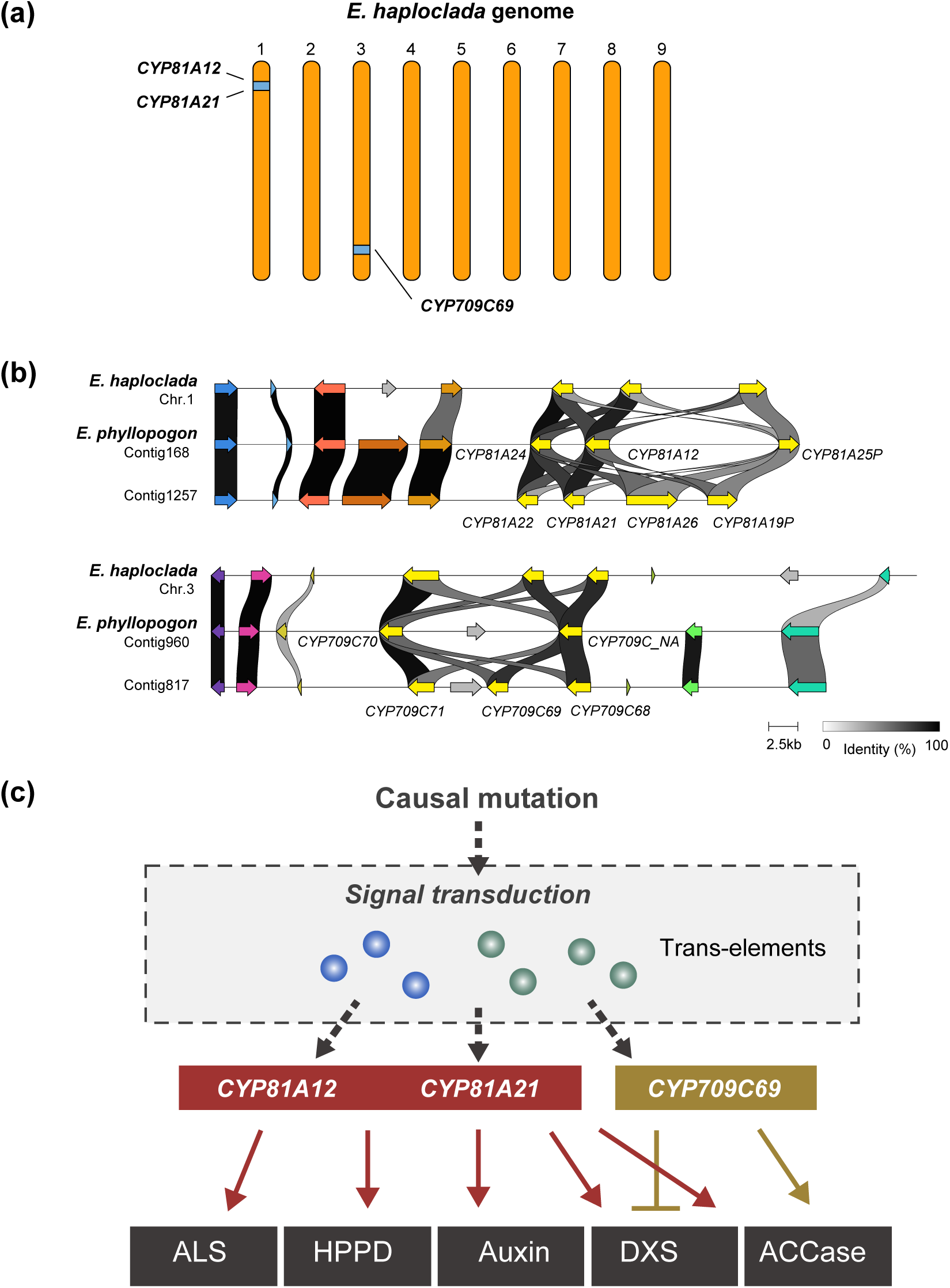
The genetic model of multiple-herbicide resistance in *Echinochloa phyllopogon*. (a) The loci of *CYP81A12*, *CYP81A21*, and *CYP709C69* orthologs in diploid *E. haploclada*. (b) The local synteny of *CYP81A* and *CYP709C* regions in the draft genome of *E. phyllopogon* (2n=4X=36) and *E. haploclada* (2n=2X=18) (Ye *et al*., 2020). (c) A working model for multiple-herbicide resistance in *E. phyllopogon*. A causal mutation activates the expression of at least three herbicide-metabolizing P450s through unknown gene-regulating systems in *E. phyllopogon*. Arrow with solid line, inactivation. T-bar, activation.

### Overexpression of the three P450 genes was observed in an MHR population found in Japan

Finally, we analyzed the generality of the overexpression of the three genes in another MHR population. Recently, a resistant population (Eoz1814) was identified from a paddy field in Niigata prefecture, Japan, the first case of herbicide resistant *E. phyllopogon* in Japan (Fig. 7a). Greenhouse experiments with commercial formulations of herbicides revealed that the resistant line purified from the population shows the decreased sensitivity to an ACCase inhibitor (cyhalofop-butyl), ALS inhibitors (pyrimisulfan, propyrisulfuron, and pinoxaden) and a synthetic auxin (quinclorac) (Fig. S8) as in the case of the MHR population found in California (Ruiz-Santaella *et al*., 2006; Chayapakdee *et al*., 2020; Dimaano *et al*., 2020). We further investigated the diclofop-methyl sensitivity of the Japanese MHR line together with an S line from Japan and MHR and S lines from California. In this assay, the Japanese MHR showed similar resistance level with the Californian MHR line (Fig. 7b). Analyses of the genes encoding herbicide target-sites, ACCase (*ACCase1* to *ACCase4*) and ALS (*ALS1* and *ALS2*), in Eoz1814 found no mutations involved in TSR mechanisms. These observations suggest that the geographically distinct two MHR populations share the resistance mechanism.

**Figure 7.**
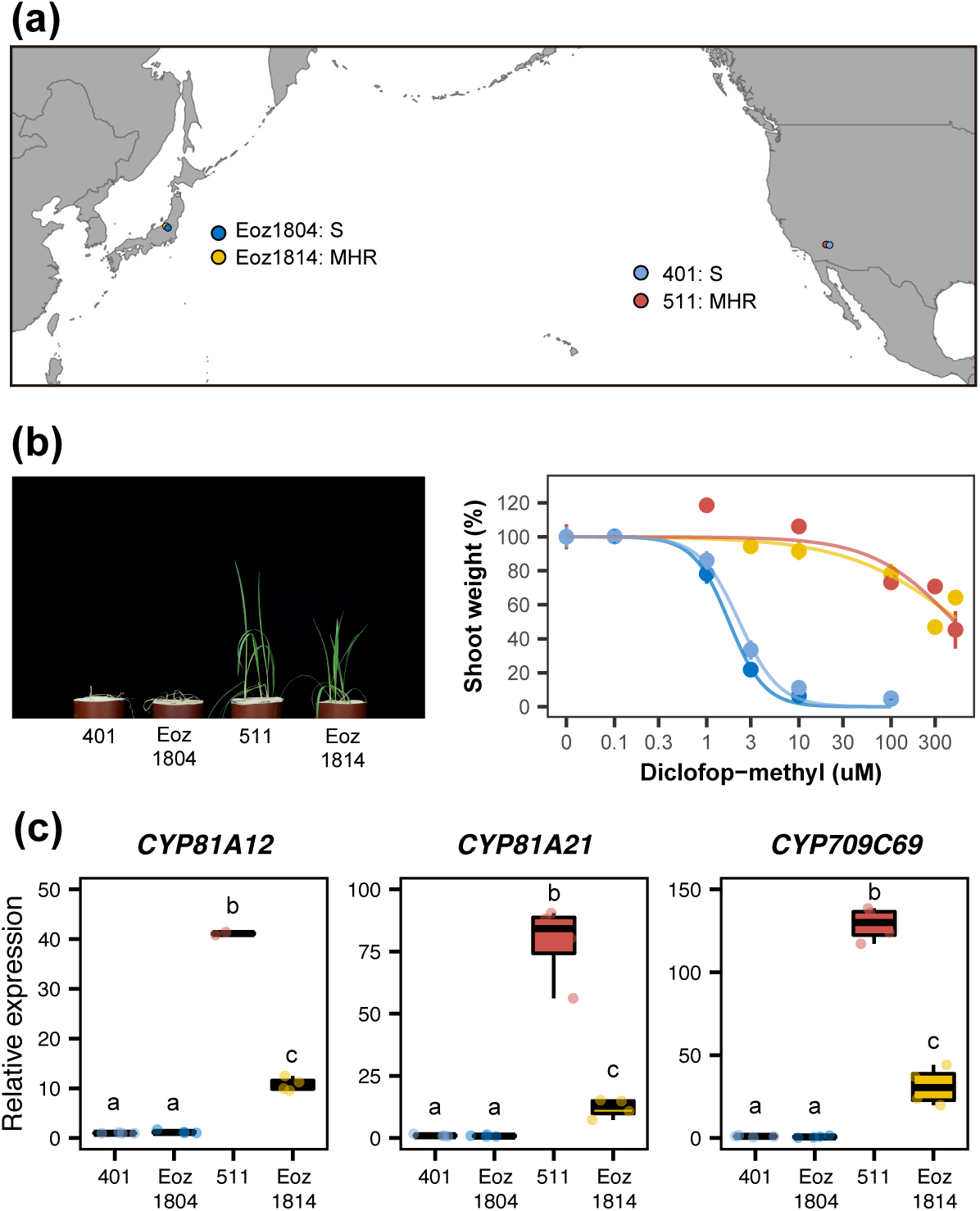
Overexpression of the three P450 genes was observed in a multiple-herbicide resistant (MHR) population found in Japan. (a) Sampling sites of the five populations. Eoz1804 and Eoz1814 are sensitive (S) and MHR, respectively. The two Californian populations (401 and 511) were used as controls. (b) Diclofop-methyl response of the four lines. The plant appearance nine days after diclofop-methyl treatment (100 μM) is shown. (c) Transcription of the three P450 genes in the shoot of 2.5-leaf stage plants. Statistically significant differences evaluated using Tukey’s HSD test are indicated by different letters (n=4).

We then analyzed the transcription of *CYP81A12*, *CYP81A21*, and *CYP706C69* by real-time PCR. The MHR line from Japan also exhibited significantly higher mRNA levels although they are less pronounced compared to the MHR line from California (Fig. 7c). The results suggest that the activation of the three P450 genes underlies the MHR in the Japanese population of *E. phyllopogon*. Meanwhile, similar diclofop-methyl resistance levels in the two MHR lines despite the difference in mRNA levels may imply the presence of additional mechanism(s) in the Japanese line.

## Discussion

Previous studies have uncovered that *E. phyllopogon* carries enzymes with broad substrate specificity, namely CYP81A12 and CYP81A21, that cause resistance to many herbicides ranging from multiple modes of action when overexpressed (Iwakami *et al*., 2019). On the other hand, the MHR *E. phyllopogon* from California exhibits resistance even to some herbicides to which the CYP81As have little or no activity, making the overexpression of CYP81As not the prime explanation to the expression of MHR trait (Dimaano *et al*., 2020, 2022). In this study, by focusing on the discrepancy between the resistance level of diclofop-methyl in the Californian MHR line and the metabolic activity of CYP81A12/21 to diclofop-methyl, we revealed that the overexpression of a novel P450 gene, *CYP709C69*, was genetically linked with MHR in all the 32 crossed progeny, where the overexpression of *CYP81A12* and *CYP81A21* has also been observed. CYP709C69 carries diclofop-acid metabolizing activity similar to CYP81A12/21 although the positions primarily hydroxylated differ between CYP709C69 and CYP81A12/21. Both hydroxylated metabolites accumulated more in the MHR line than the S line, which is most likely caused by the overexpressed CYP709C69 and CYP81A12/21. Thus, the cooperative action of the three P450s on the active form of the herbicide underlies the high-level resistance to diclofop-methyl resistance in the MHR line.

Unlike the broad substrate specificity of CYP81A12/21, CYP709C69 only metabolized clomazone among the 46 herbicides that were further tested: *A. thaliana* expressing this gene exhibited higher sensitivity to clomazone. Regardless of the high substrate specificity, clomazone metabolism in particular needs to be further investigated. Firstly, the study will accumulate molecular insight about plant genes for herbicide activation, where only a single gene to activate ACCase herbicides was reported (Cummins & Edwards, 2004). Secondly, the role of herbicide-activating enzymes is important in the context of herbicide-resistance evolution. Reduced herbicide-activation is associated with triallate resistance in *Avena fatua* found in Montana (Kern *et al*., 1996), which can be caused by the disruption of a gene with similar function of CYP709C69. It is interesting to investigate whether clomazone resistance can be evolved by knocking out *CYP709C69*. Lastly, the increased clomazone sensitivity in *A. thaliana* implies that resistant weeds with enhanced herbicide metabolism may exhibit negative cross-resistance to some herbicides depending on the properties of the causal enzymes. The insight further highlights the importance of the findings. However, it is of note that the MHR line from California is resistant to clomazone (Yasuor *et al*., 2008) despite the overexpression of CYP709C69. This is likely explained by the clomazone metabolizing activity of CYP81A12/21. Our previous studies showed that *A. thaliana* expressing *CYP81A12* or *CYP81A21* became resistant to clomazone (Guo *et al*., 2019) and that they oxidize clomazone to an unidentified metabolite (Dimaano *et al*., 2020). The higher expression of *CYP81A12/21* than *CYP709C69* in the MHR line (Fig. 2c) suggests that inactivating forces (CYP81A12/21) are the greater than activating forces (CYP709C69).

The present study represents a major step forward toward a complete understanding of the plant metabolism of the “model herbicide” diclofop-methyl, which is one of the well-understood herbicides in biochemistry in planta. We disclosed the genes producing the two of the three hydroxylated diclofop-acid metabolites found in wheat in the early 90’s (Tanaka *et al*., 1990; Zimmerlin & Durst, 1992). CYP81As and CYP709Cs in both *E. phyllopogon* and rice produced M1 and M2, respectively, suggesting the conserved function in other species. The genes in wheat/barley, major crops for diclofop-methyl usage, and in *L. rigidum*, a major weed well-known for the metabolism-based resistance to diclofop-methyl, are worthy of investigation. In our experiments, we observed that yeast expressing CYP81A (possibly CYP709C as well) produced a small peak of M3, which likely corresponds to a metabolite with the 2,3-dichloro-4-hydroxyphenoxy moiety detected in wheat (Fig. 3c). The reason why M3 was not detected in *E. phyllopogon* is unclear, but this could be due to differences in the experimental systems between yeast and *E. phyllopogon*: In the yeast whole cell assay, the cells were treated with a 300 μM diclofop-methyl solution for 24 h, while *E. phyllopogon* was treated with a 30 μM diclofop-methyl solution for 30 min. Thus, the yeast system subjected much higher concentration of the substrate to the enzymes, possibly forcing a metabolic reaction to produce M3. It is also possible that M3 is more susceptible to further metabolization in *E. phyllopogon* than M1 and M2, reducing the actual amount of M3 and M3-glucoside than those of M1 and M2 in *E. phyllopogon*. Analysis of other species in addition to the characterization of the M3 metabolite will provide insight into these possibilities.

Trough the detailed diclofop-methyl metabolism study, we disclosed that CYP81As mediate chlorine migration reaction in the process of diclofop hydroxylation. The hydroxylation of the aromatic ring can cause the migration of the original functional group such as halogen, alkyl group, and carboxyl group, to the adjacent carbon, which is called NIH-shift (Zhao *et al*., 2018). P450s are among the major enzymes involved in NIH-shift although the specific P450 members that cause NIH-shift are rarely reported. 2,4-D and diclofop-methyl are the two major herbicides involving NIH-shift in their metabolism *in planta* (Dimaano & Iwakami, 2021). The biological influence of the NIH-shift reactions in these herbicides is not well-understood. The reactions may influence the accessibility of the enzymes governing the following reactions of the metabolites, which may further influence plant herbicide sensitivity.

This study provides another aspect in the mechanism of MHR in agricultural weeds: MHR can be caused by the result of genetically-linked simultaneous activation of genes encoding herbicide-detoxifying enzymes with different enzymatic properties. The idea differs from a mere stepwise accumulation of multiple mechanisms to broaden the spectrum of the resistance as suggested in other species (e.g. Preston *et al*., 1996; Giacomini *et al*., 2020). Similar concept is suggested in the mode of action of herbicide safener, chemical agents that reduce the phytotoxicity of herbicides to crop plants, where a concerted upregulation of multiple herbicide-metabolism related genes has been proposed as the mechanism (Riechers *et al*., 2010) although no clear genetic investigations for the concerted upregulation has been provided. Thus, it is possible that the safener-mediated concerted regulations are caused by the stimulations to the multiple independent gene regulation networks, which is clearly distinct from the concept of genetically-linked simultaneous activation of herbicide metabolizing genes provided here. Intriguingly, we observed the co-segregations of overexpression and MHR in at least three more P450 genes in the F6 lines, i.e., *CYP72A122*, *CYP72A252v1*, and *CYP72A252v2* (Fig. 2e). If any of the genes carry herbicide-metabolizing function, the scope of resistance of the MHR line would be expanded.

The three P450 overexpression may correspond to a constitutive activation of some plant defense against biotic or abiotic stress, and to the expression of one or more natural compound pathway(s). CYP81As in maize are involved in the biosynthesis of zealexins, antimicrobial substances (Ding *et al*., 2020), where a cluster of genes including *CYP81A* and other P450 genes are activated in response to pathogen infestation, leading to zealexin biosynthesis. Given a similar metabolic regulation system is conserved in *E. phyllopogon*, the overexpression of the three P450 genes may be a result of constitutive overexpression of plant defense system. Elucidation of the regulatory system and the endogenous function of each P450 may provide insight into the fitness costs associated with resistance expression and hints for the control of MHR plants that are hardly controlled by herbicide chemistries.

Another interesting finding is that overexpression of the three P450s was also found in the Japanese line. It is unlikely that herbicide selection pressure acted on *CYP709C69*, since diclofop and clomazone, which were the only herbicides that CYP709C69 metabolized in our study, have never been registered in rice cropping in Japan. Thus, it is more likely that other selection pressures, such as ALS inhibitors, have selected for a mutation that activate the regulatory system for expression of genes such as *CYP81A12* and *CYP81A21* that metabolize these herbicides. The fact that the Japanese line has the same diclofop-methyl resistance trait as the MHR line from California, even though they were selected not by diclofop-methyl, supports the idea that the gene regulatory system proposed in the Californian line is a common endogenous system shared by other *E. phyllopogon*. In herbicide resistance, molecular convergence is frequently observed in the case of target-site resistance (Baucom, 2019). On the other hand, the convergence in metabolic resistance remains poorly understood. Elucidation of the causal mechanisms that confer resistance in these MHR *E. phyllopogon* is expected to provide new insights in understanding the mechanisms of stress tolerance evolution in plants.

## Acknowledgements

We thank Dr. David Nelson for naming the P450s, Dr. Niña Gracel Dimaano for English critique, and Yuuri Nakamura for technical assistance. RNA-seq was supported by the NODAI Genome Research Center, Tokyo University of Agriculture under Grant Cooperative Research of the Genome Research for Bio-Resource. Computations were partially performed on the NIG supercomputer at ROIS National Institute of Genetics. This work was supported by a JSPS KAKENHI grant no. 15H06072, 17K15234, 19H02955 (to S.I.), and in part by the Program for the Development of Next-generation Leading Scientists with Global Insight (L-INSIGHT), sponsored by the Ministry of Education, Culture, Sports, Science and Technology (MEXT), Japan.

## Author contributions

Conceptualization, S.I.; Research design, S.I., H.S., M.M., Y.Y., T.Y., K.T., S.T., T.T.; RNA-sequencing, K.T.; Bioinformatics, S.I., H.S., T.K.; LC-MS/MS and NMR, H.S., Y.Y., M.M.; Japanese line characterization, K.T., A.U., S.A., M.H.; Gene characterization, H.S., S.I., T.Y.; Writing, S.I., H.S., M.M.; Visualization, S.I., H.S., T.K.; Supervision, S.I., T.T.; Funding acquisition, S.I.

## Data availability

The mRNA-Seq data have been deposited in the DDBJ Sequence Read Archive (DRA) database (accession number DRA013092).

## Supplementary data

**Table S1. Primers used in this study**

a, Primer concentration in real-time PCR. b, Developed in Tanigaki *et al*., (2021). c, Developed in Czechowski *et al*., (2005). d, Developed in Iwakami *et al*., (2014b).

**Table S2. The summary of herbicides used for the functional characterizations of *CYP709C69***

The herbicide classifications follow the HRAC system (https://www.hracglobal.com/). Another mode of action was annotated for quinclorac (cellulose synthesis) and bromoxynil (uncouplers).

**Figure S1. Annotation correction**

The original annotations for Contig817_pilon.380 (a) and Contig222_pilon.21 (b) were shown as boxes with solid lines. The missing exons in the original annotations (Ye *et al*., 2020) were shown as boxes with dotted lines. The new gene models were distinguished with the suffix of a and b. The comparison of Contig222_pilon.21.a and Contig222_pilon.21.b suggests that the latter carries a single nucleotide deletion in the first exon at the position of an asterisk (*), leading to a premature stop codon.

**Figure S2. Detection of differentially expression genes by RNA-seq**

(a) Multi-dimensional scaling (MDS) plot of the RNA-seq libraries of the sensitive (S) and multiple-herbicide resistant (MHR) lines. (b) Volcano plot of the S and MHR lines. Differentially expressed genes (fold-change>3, FDR<0.05) are shown in dark gray. Upregulated P450 genes (Iwakami *et al*., 2014a)are shown in yellow.

**Figure S3. Diclofop-methyl sensitivity of P450 genes in *Echinochloa phyllopogon***

Independent calli were grown on the media with diclofop-methyl for three weeks. (a) Candidate P450 genes. (b) CYP709C genes.

**Figure S4. Diclofop-methyl sensitivity of *Arabidopsis thaliana* transformed with P450 genes in *Echinochloa phyllopogon***

(a) Transcript levels of T3 homozygous lines generated in this study. (b) Response to 30 μM diclofop-methyl. Bars represent SEM. *CYP81A21*#6-1 was previously established line (Iwakami *et al*., 2014a). The line exhibited the highest mRNA level among the tested lines, and showed resistance to many herbicides (Iwakami *et al*., 2014a; Dimaano *et al*., 2020).

**Figure S5. ^1^H-NMR spectra for OH-diclofop-acid of M1 (A) and M2 (B)**

Asterisks indicate the signals related to impurities.

**Figure S6. Herbicide metabolizing activity of CYP709C69**

(a) Rice calli responses to herbicides inhibiting acetyl-CoA carboxylase. Independent calli were grown on the media with herbicides for three weeks. (b) *Arabidopsis thaliana* responses to herbicides. Two highly expressing lines (#31-5 and #17-2) (Fig. S4) were grown on the media with herbicides for 12 days. The herbicide classifications follow the HRAC system (https://www.hracglobal.com/).

**Figure S7. Progeny association of the mRNA levels of *CYP709C69***

Shoot mRNA levels (four plants in bulk) were quantified by real-time PCR in 16 F6 lines that were previously characterized (Iwakami *et al*., 2019; Chayapakdee *et al*., 2020). R6-R/S is a segregating line.

**Figure S8. Herbicide sensitivity of *Echinochloa phyllopogon* in Japan**

(a) Herbicides and the growth stage of *E. phyllopogon* used for the assay. 1x and 1/3x field rate was applied. The 1x rate of quinclorac was followed by that registered in the USA (Yasuor *et al*., 2012). (b) Herbicide response of resistant *E. phyllopogon* (Eoz1814) found in Japan. Shoot dry weight of 13 days after herbicide applications were evaluated. Bars, SEM (n=5). Line 401, originated from California, was used as a sensitive control.

