## Supplementary material for "Genetically-linked simultaneous overexpression of multiple herbicide-metabolizing genes for broad-spectrum resistance in an agricultural weed *Echinochloa phyllopogon*": Fig. S1

(a)

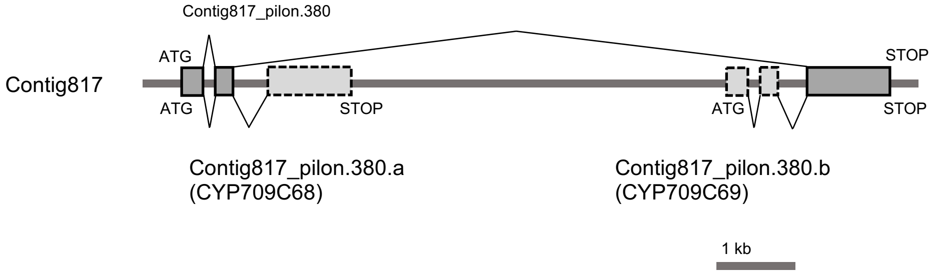


(b)

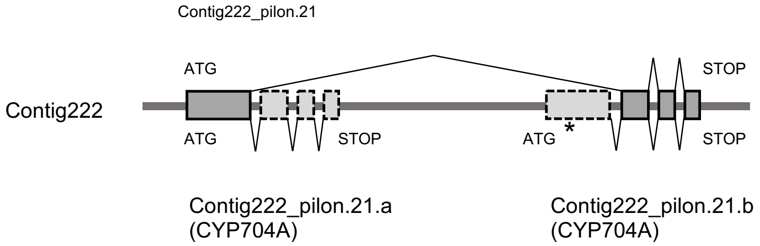


**Figure S1. Annotation correction**

The original annotations for Contig817_pilon.380 (a) and Contig222_pilon.21 (b) were shown as boxes with solid lines. The missing exons in the original annotations [18] were shown as boxes with dotted lines. The new gene models were distinguished with the suffix of a and b. The comparison of Contig222_pilon.21.a and Contig222_pilon.21.b suggests that the latter carries a single nucleotide deletion in the first exon at the position of an asterisk (*), leading to a premature stop codon.
