## Supplementary material for "Genetically-linked simultaneous overexpression of multiple herbicide-metabolizing genes for broad-spectrum resistance in an agricultural weed *Echinochloa phyllopogon*": Fig. S2

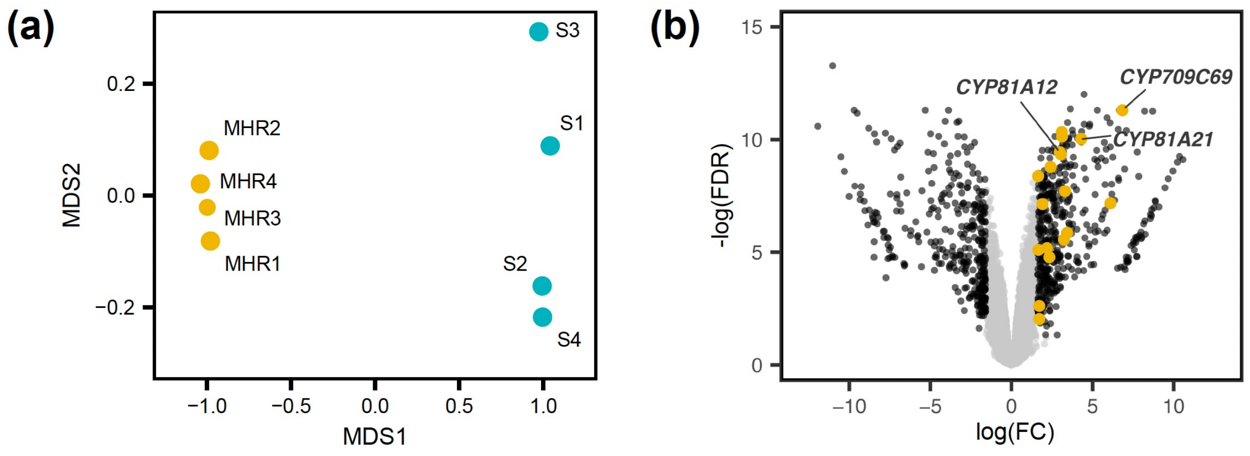


**Figure S2. Detection of differentially expression genes by RNA-seq**
(a) Multi-dimensional scaling (MDS) plot of the RNA-seq libraries of the sensitive (S) and multiple-herbicide resistant (MHR) lines. (b) Volcano plot of the S and MHR lines. Differentially expressed genes (fold-change>3, FDR<0.05) are shown in dark gray. Upregulated P450 genes (Iwakami *et al.*, 2014a)are shown in yellow.
