## Supplementary material for "Genetically-linked simultaneous overexpression of multiple herbicide-metabolizing genes for broad-spectrum resistance in an agricultural weed *Echinochloa phyllopogon*": Fig. S3

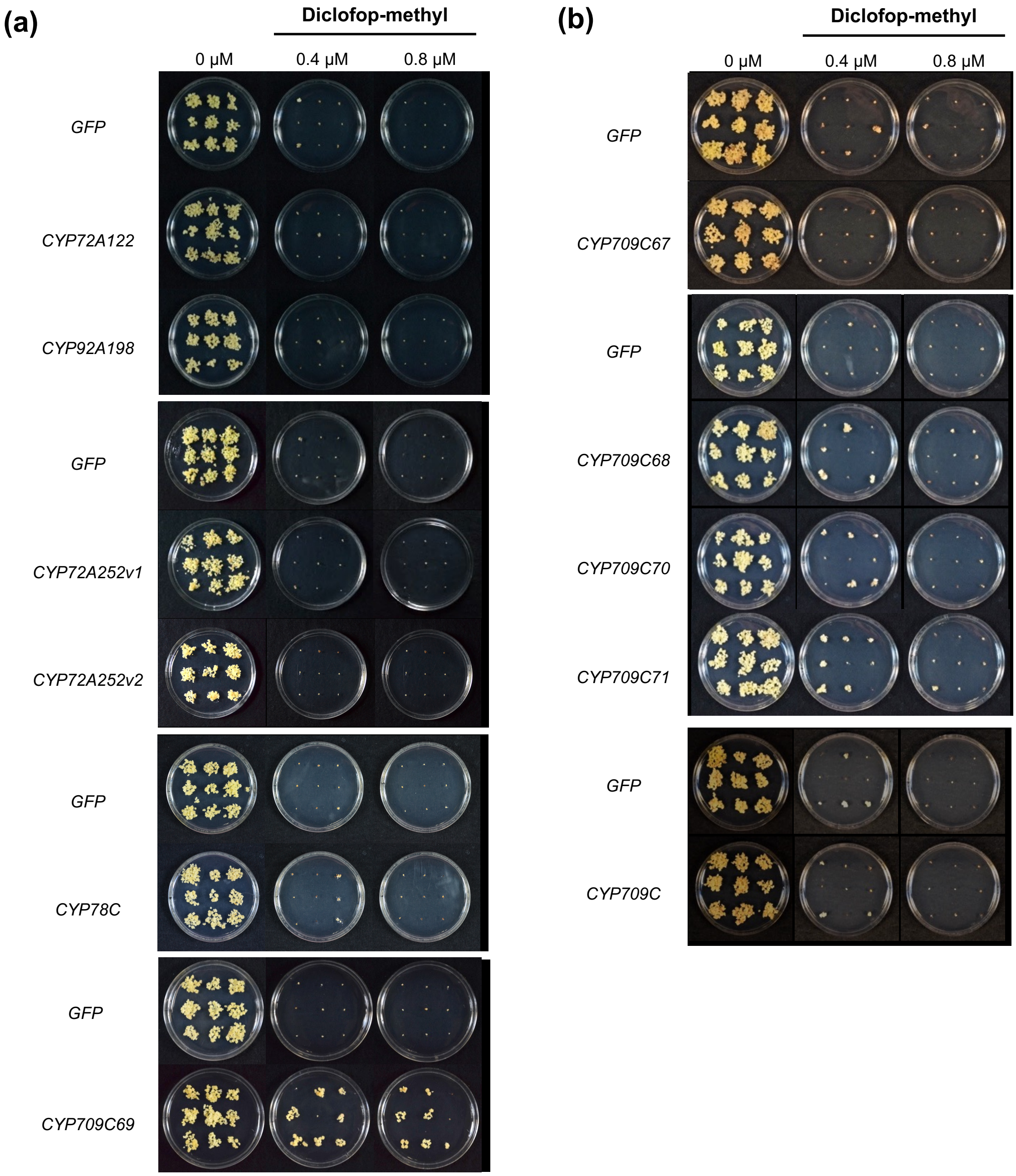


**Figure S3. Diclofop-methyl sensitivity of P450 genes in *Echinochloa phyllopogon***
Independent calli were grown on the media with diclofop-methyl for three weeks.
(a) Candidate P450 genes. (b) CYP709C genes.
