## Supplementary material for "Genetically-linked simultaneous overexpression of multiple herbicide-metabolizing genes for broad-spectrum resistance in an agricultural weed *Echinochloa phyllopogon*": Fig. S4

(a) (b)

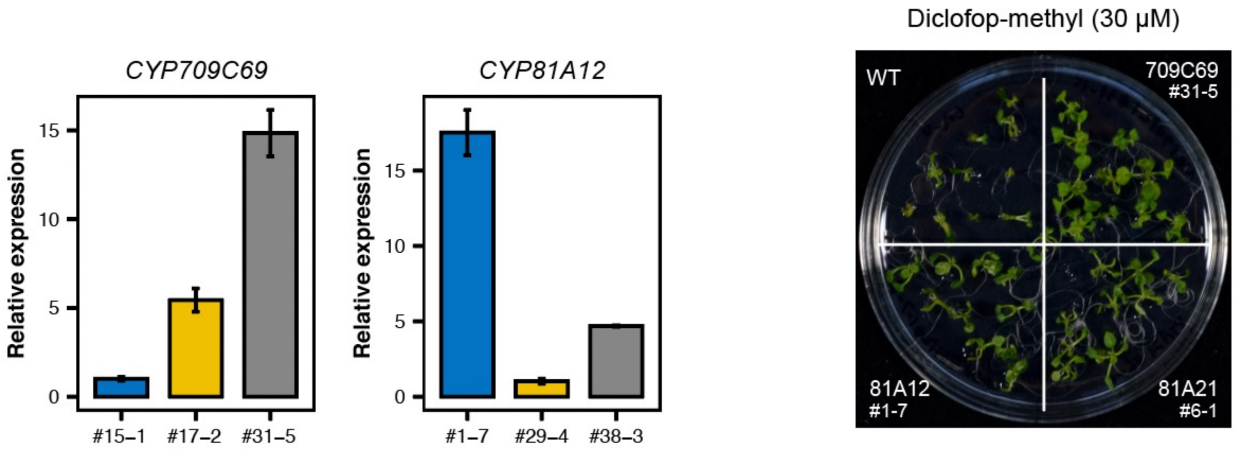


**Figure S4. Diclofop-methyl sensitivity of *Arabidopsis thaliana* transformed with P450 genes in *Echinochloa phyllopogon***

(a) Transcript levels of T3 homozygous lines generated in this study. (b) Response to 30 μM diclofop-methyl. Bars represent SEM. *CYP81A21*#6-1 was previously established line (Iwakami *et al.*, 2014a). The line exhibited the highest mRNA level among the tested lines, and showed resistance to many herbicides (Iwakami *et al.*, 2014a; Dimaano *et al.*, 2020).
