## Supplementary material for "Genetically-linked simultaneous overexpression of multiple herbicide-metabolizing genes for broad-spectrum resistance in an agricultural weed *Echinochloa phyllopogon*": Fig. S5

(a)

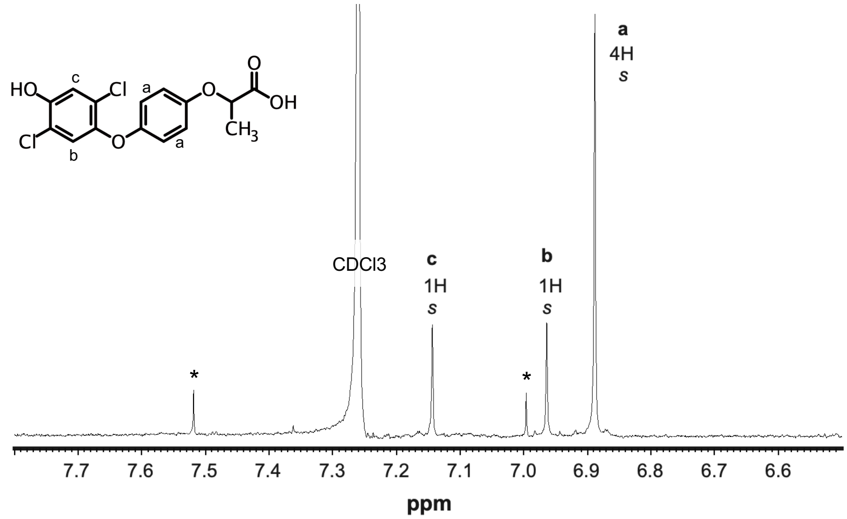


(b)

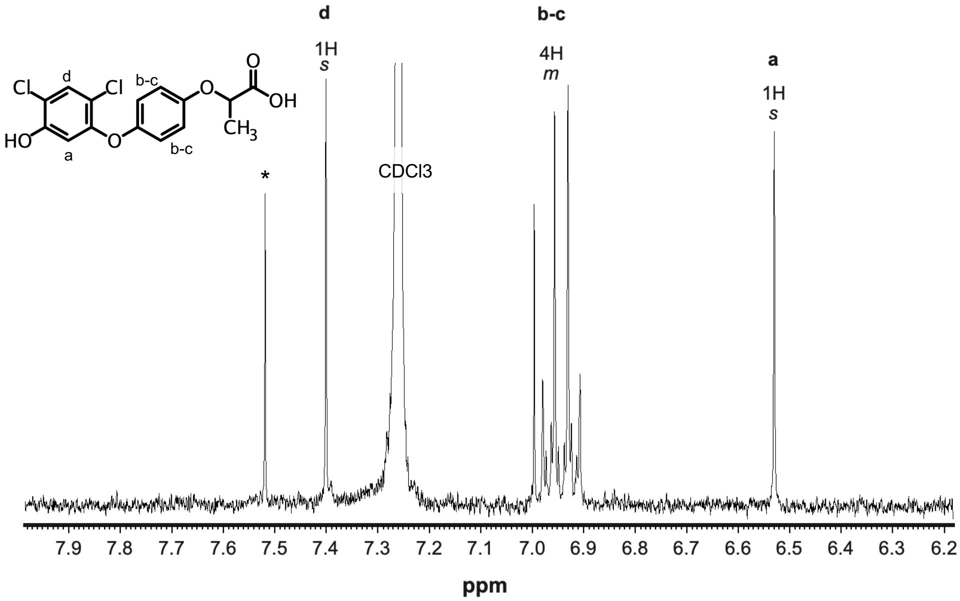


**Figure S5. [1]H-NMR spectra for OH-diclofop-acid of M1 (a) and M2 (b)**
Asterisks indicate the signals related to impurities.
