## Supplementary material for "Genetically-linked simultaneous overexpression of multiple herbicide-metabolizing genes for broad-spectrum resistance in an agricultural weed *Echinochloa phyllopogon*": Fig. S6

**(a)**

### Group 1: Inhibition of Acetyl CoA Carboxylase

***Aryloxphenoxy-propionates***

**Fenoxaprop-P-ethyl** | **Metamifop** | **Clodinafop-propargyl**

0 μM | 0.1 μM | 0.3 μM | 0.3 μM | 1 μM | 3 μM

*GFP*

*CYP709C69*

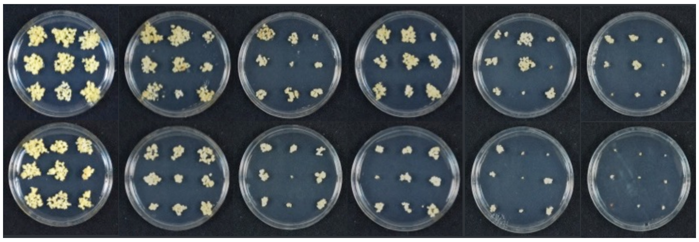

0 nM | 40 nM | 80 nM | 120 nM

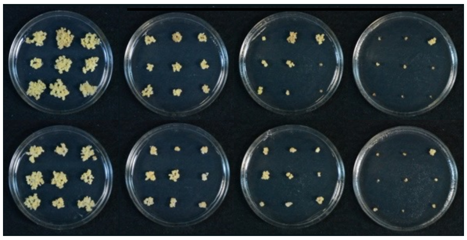

**Cyhalofop-buthyl**

0 μM | 1 μM | 3 μM | 10 μM

*GFP*

*CYP709C69*

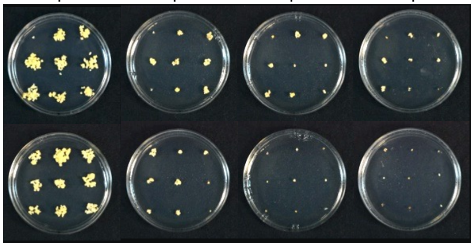

**Quizalofop-ethyl**

0 μM | 2 μM | 4 μM

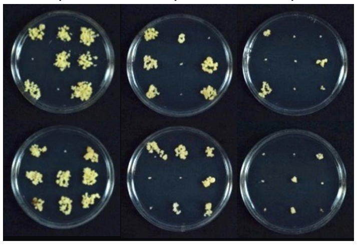

***Cyclohexanediones***

**Tralkoxydim**

0 μM | 0.3 μM | 0.6 μM

*GFP*

*CYP709C69*

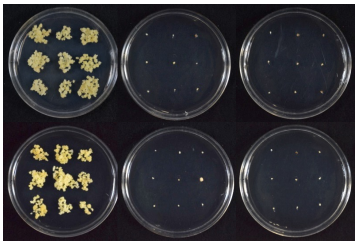

***Not annotated***

**Pinoxaden**

0 μM | 0.05 μM | 0.1 μM

*GFP*

*CYP709C69*

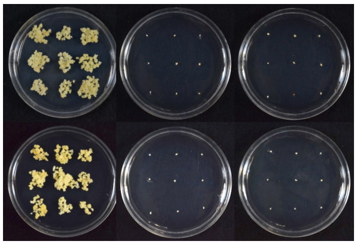

(b)

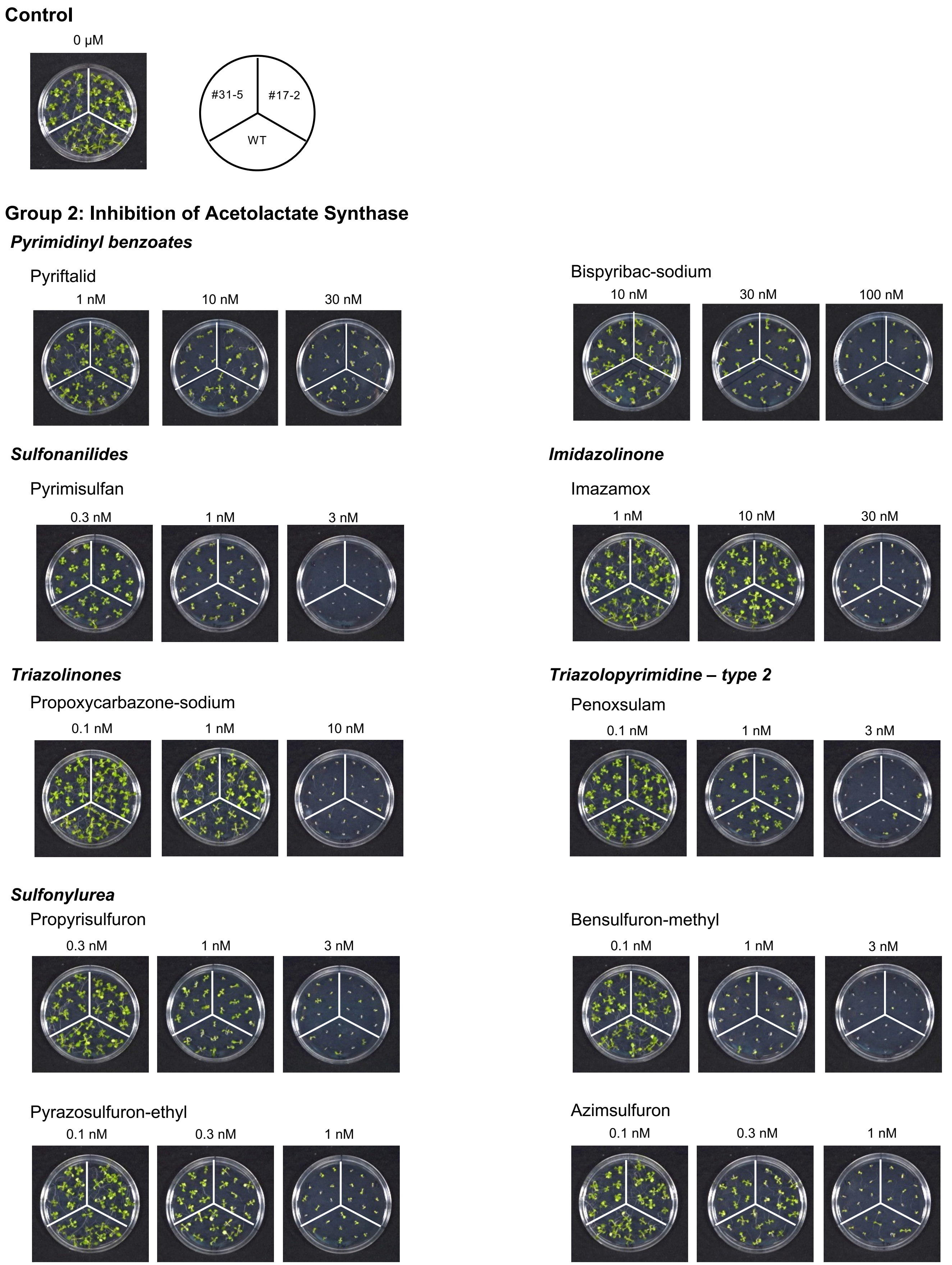
Control
0 µM
#31-5
#17-2
WT
Group 2: Inhibition of Acetolactate Synthase
Pyrimidinyl benzoates
Pyriftalid
1 nM
10 nM
30 nM
Bispyribac-sodium
10 nM
30 nM
100 nM
Sulfonanilides
Pyrimisulfan
0.3 nM
1 nM
3 nM
Imidazolinone
Imazamox
1 nM
10 nM
30 nM
Triazolinones
Propoxycarbazone-sodium
0.1 nM
1 nM
10 nM
Triazolopyrimidine – type 2
Penoxsulam
0.1 nM
1 nM
3 nM
Sulfonylurea
Propyrisulfuron
0.3 nM
1 nM
3 nM
Bensulfuron-methyl
0.1 nM
1 nM
3 nM
Pyrazosulfuron-ethyl
0.1 nM
0.3 nM
1 nM
Azimsulfuron
0.1 nM
0.3 nM
1 nM

Chlorsulfuron

1 nM | 3 nM | 10 nM

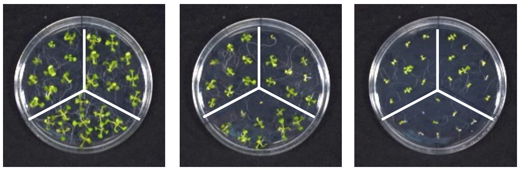

### Group 3: Inhibition of Microtubule Assembly

#### *Dinitroanilines*

Trifluralin

1 μM | 3 μM | 10 μM

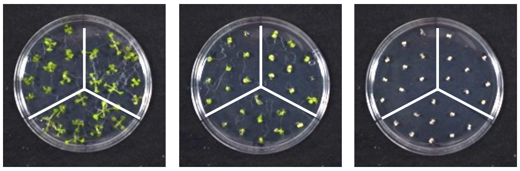

### Group 4: Auxin Mimics

#### *Phenoxy-carboxylates*

2,4-D

30 nM | 100 nM | 300 nM

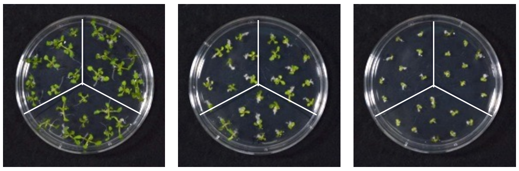

#### *Quinoline-carboxylates*

Quinclorac

10 μM | 60 μM | 300 μM

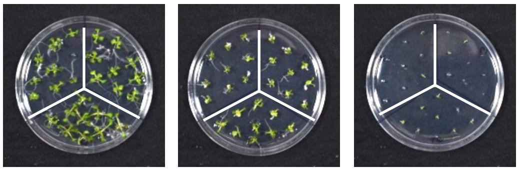

### Group 5: $D_1$ Serine 264 binders (and other non-histidine 215 binders)

#### *Amides*

Propanil

1 μM | 10 μM | 100 μM

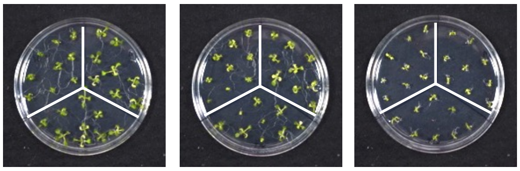

#### *Triazines*

Cyanazine

0.3 μM | 1 μM | 3 μM

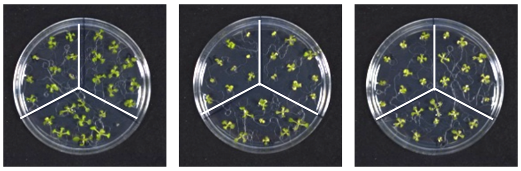

### Group 6: $D_1$ Histidine 215 binders

#### *Not annotated*

Bentazone

1 μM | 3 μM | 30 μM

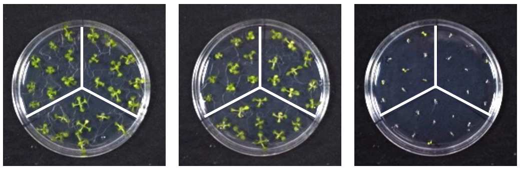

#### *Nitriles*

Bromoxynil

1 μM | 3 μM | 10 μM

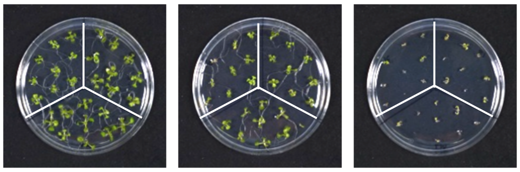

### Group 9: Inhibition of Enolpyruvyl Shikimate Phosphate Synthase

#### *Not annotated*

Glyphosate

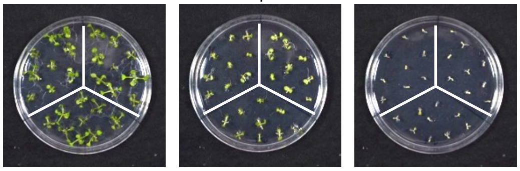


### Group 10: Glutamine Synthetase

#### *Phosphinic acids*

Glufonsinate

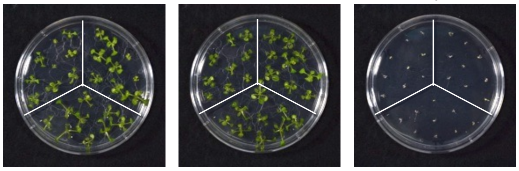


### Group 12: Inhibition of Phytoene Desaturase

#### *Pheyl-ethers*

Diflufenican

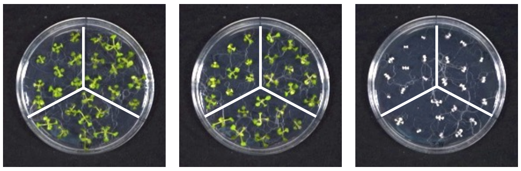


#### *Diphenyl heterocycles*

Fluridone

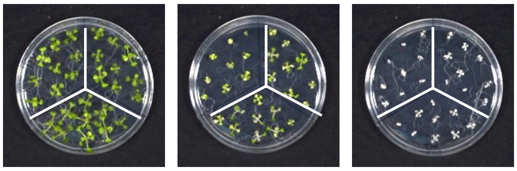


#### *N-Phenyl heterocycles*

Norflurazon

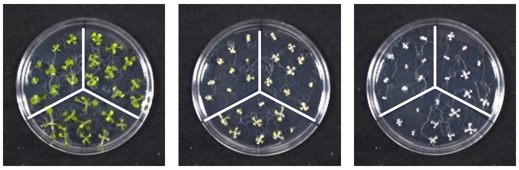


### Group 13: Inhibition of Deoxy-D-Xyulose Phosphate Synthase

#### *Isoxazolidinones*

Clomazone

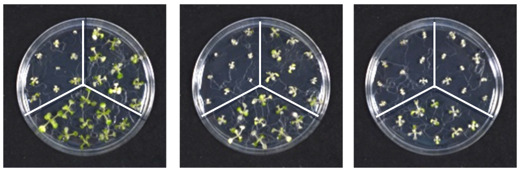

### Group 14: Inhibition of Protoporphyrinogen Oxidase

#### *Not annotated*

Pyraclonil

0.3 nM 1 nM 3 nM

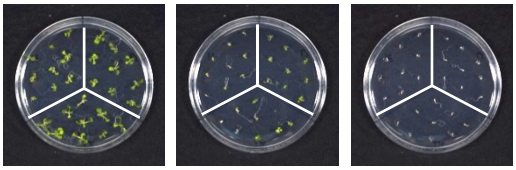

#### *N-Phenyl-oxadiazolones*

Oxadiargyl

100 nM 300 nM 600 nM

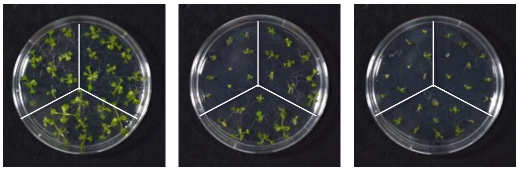

#### *N-Phenyl-imides*

Flumioxazin

0.1 nM 1 nM 10 nM

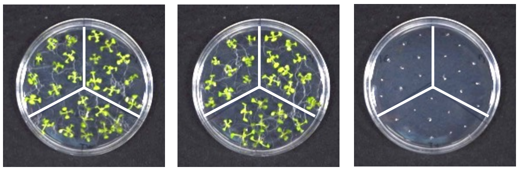

### Group 15: Inhibition of Very Long-Chain Fatty Acid Synthesis

#### *α-Chloroacetamides*

Butachlor

3 μM 6 μM 20 μM

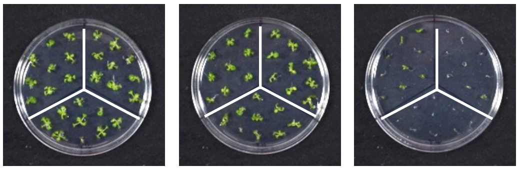

Pretilachlor

3 μM 10 μM 30 μM

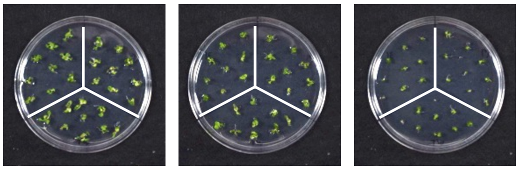

#### *Thiocarbamates*

Molinate

60 μM 100 μM 300 μM

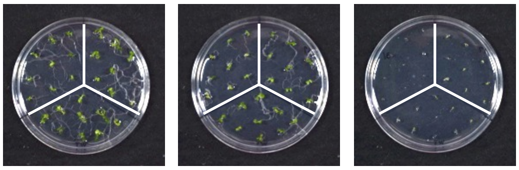

Thiobencarb

10 μM 30 μM 100 μM

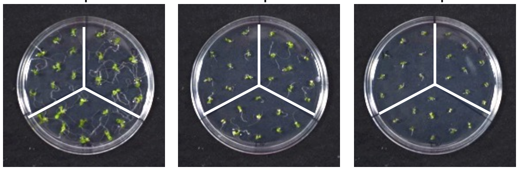

#### *Isoxazolines*

Fenoxasulfone

1 μM 100 μM 300 μM

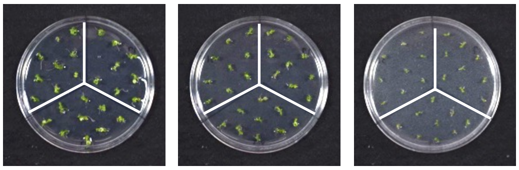

Pyroxasulfone

1 μM 10 μM 30 μM

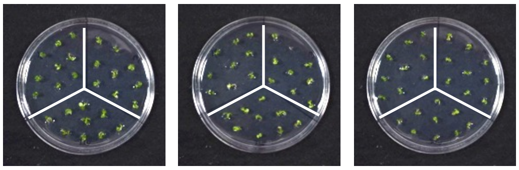

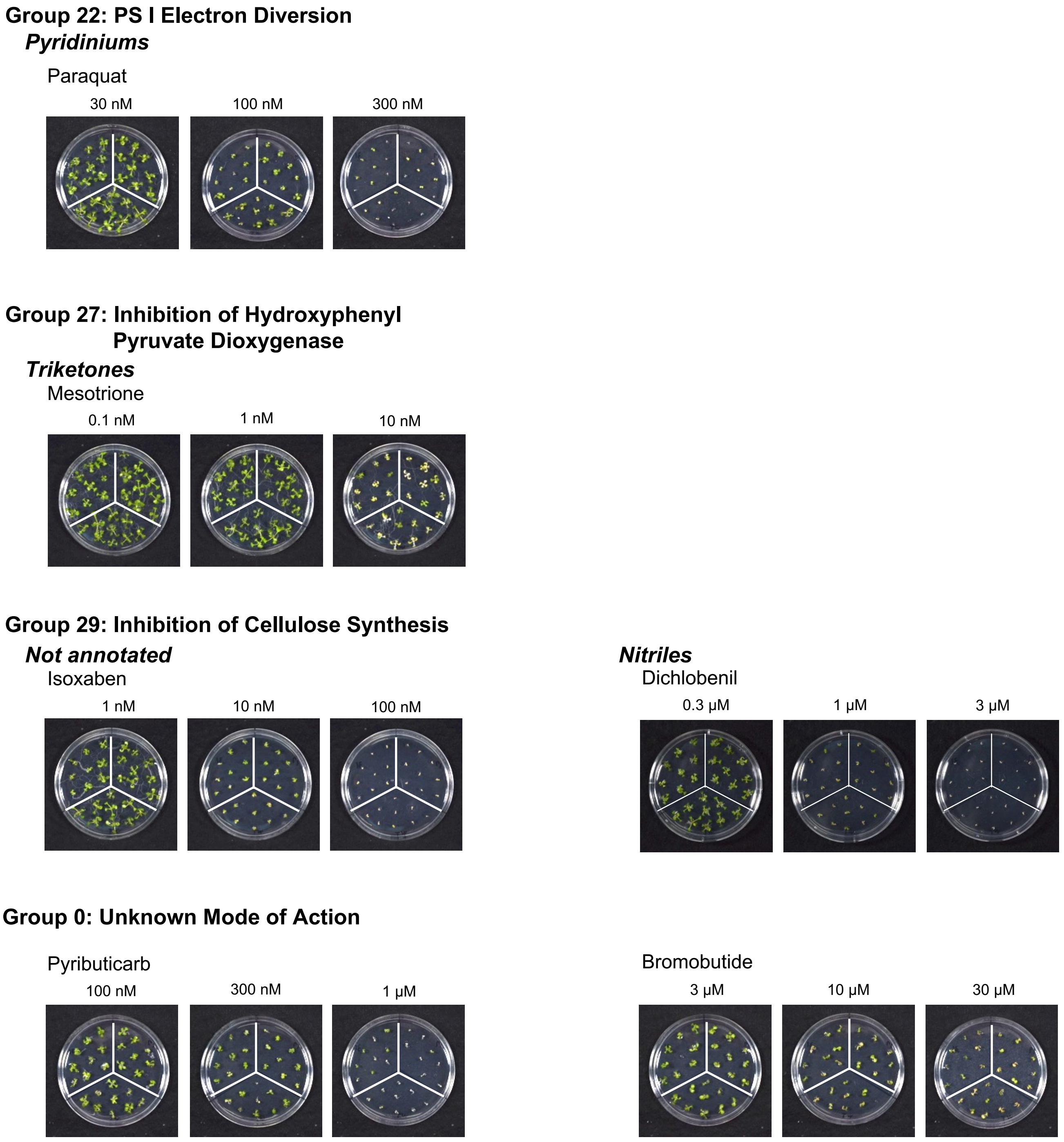


**Figure S6. Herbicide metabolizing activity of CYP709C69**

(a) Rice calli responses to herbicides inhibiting acetyl-CoA carboxylase. Independent calli were grown on the media with herbicides for three weeks. (b) *Arabidopsis thaliana* responses to herbicides. Two highly expressing lines (#31-5 and #17-2) (Fig. S4) were grown on the media with herbicides for 12 days. The herbicide classifications follow the HRAC system (https://www.hracglobal.com/).
