## Supplementary material for "Genetically-linked simultaneous overexpression of multiple herbicide-metabolizing genes for broad-spectrum resistance in an agricultural weed *Echinochloa phyllopogon*": Fig. S7

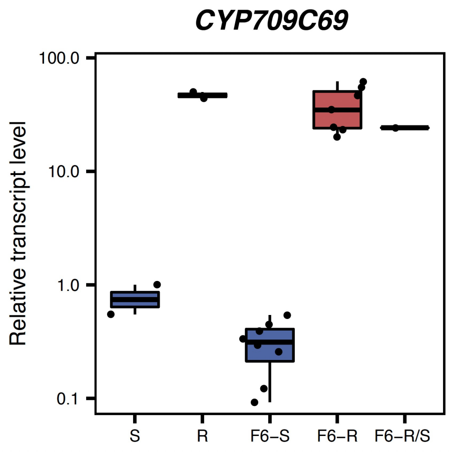


**Figure S7. Progeny association of the mRNA levels of *CYP709C69***
Shoot mRNA levels (four plants in bulk) were quantified by real-time PCR in 16 F6 lines that were previously characterized (Iwakami *et al.*, 2019; Chayapakdee *et al.*, 2020). R6-R/S is a segregating line.
