## Supplementary material for "Genetically-linked simultaneous overexpression of multiple herbicide-metabolizing genes for broad-spectrum resistance in an agricultural weed *Echinochloa phyllopogon*": Fig. S8

**(a)**

| Mode of action | Herbicide | Trade name | Company | 1x rate (g a.i./ha) | Applied leaf-sage |
|---|---|---|---|---|---|
| ACCase | Cyhalfop-butyl | Clincher EW | Dow AgroSciences, Tokyo, Japan | 300 | 4.5 |
| ALS | Pyrimisulfan | Best Partner | Kumiai chemical industry, Tokyo, Japan | 67 | 2.5-3 |
| ALS | Propyrisulfuron | ZETA-ONE | Sumitomo Chemical Company, Tokyo, Japan | 90 | 2.5-3 |
| ALS | Penoxsulam | Wide Attack SC | Dow AgroSciences, Tokyo, Japan | 37.5 | 4.5 |
| Auxin | Quinclorac | Quinclorac 250 SC | BASF, Ludwigshafen, Germany | 560 | 2.5-3 |

**(b)**

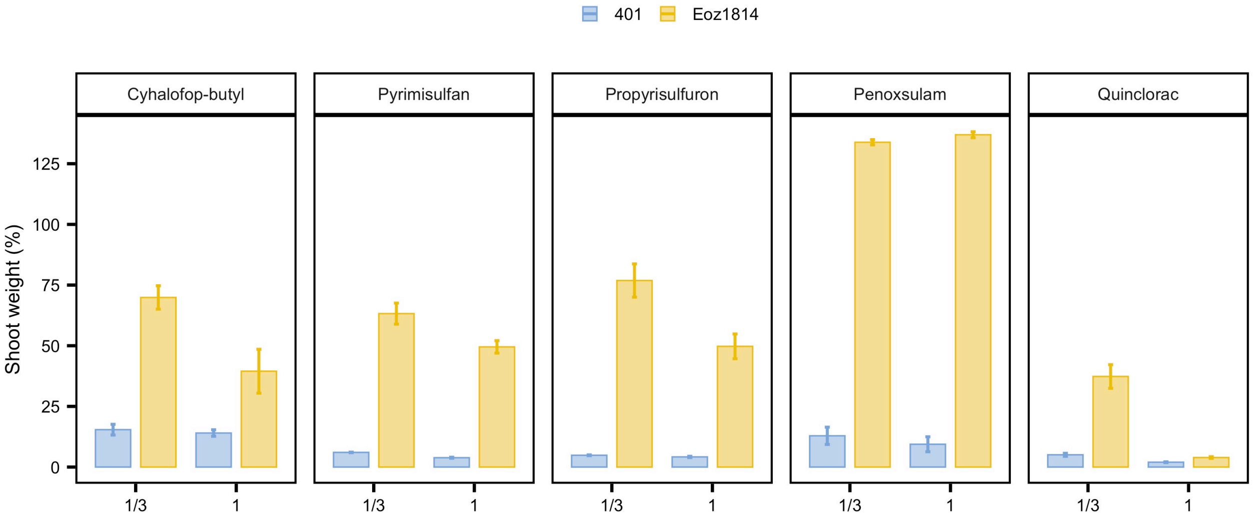


**Figure S8. Herbicide sensitivity of *Echinochloa phyllopogon* in Japan**
(a) Herbicides and the growth stage of *E. phyllopogon* used for the assay. 1x and 1/3x field rate was applied. The 1x rate of quinclorac was followed by that registered in the USA (Yasuor *et al.*, 2012). (b) Herbicide response of resistant *E. phyllopogon* (Eoz1814) found in Japan. Shoot dry weight of 13 days after herbicide applications were evaluated. Bars, SEM (n=5). Line 401, originated from California, was used as a sensitive control.
