## Supplementary material for "Genetically-linked simultaneous overexpression of multiple herbicide-metabolizing genes for broad-spectrum resistance in an agricultural weed *Echinochloa phyllopogon*": Table S1

### Table S1. Primers used in this study.

| Gene | Species | Forward Primer (5'-3') | Reverse Primer (5'-3') | Concentration[a] |
|---|---|---|---|---|
| **pCAMBIA cloning** | | | | |
| *CYP709C67* | *E. phyllopogon* | TACAATTACAGTCGAATGGGCGTCGCCGGG | CCGCTTTACTTGTACCTATGGCTCCAAGAG | |
| *CYP709C68* | *E. phyllopogon* | TACAATTACAGTCGAATGGGCGTGGCATG | CCGCTTTACTTGTACTTACACCTCCAAAC | |
| *CYP709C69* | *E. phyllopogon* | TACAATTACAGTCGAATGGGTCTGGCTTGGAT | CCGCTTTACTTGTACTTACACCTCCACGCG | |
| *CYP709C70* | *E. phyllopogon* | TACAATTACAGTCGAATGGGTTACGGGTGG | CCGCTTTACTTGTACCTACGCGTCGTCAGCAC | |
| *CYP709C71* | *E. phyllopogon* | TACAATTACAGTCGAATGGGGTACGGGTGG | CCGCTTTACTTGTACCTACGCGTCGTCAGCAC | |
| *CYP709C* | *E. phyllopogon* | TACAATTACAGTCGAATGGTGGACATGGGC | CCGCTTTACTTGTACTTAATATTCTCCCA | |
| *CYP72A122* | *E. phyllopogon* | TACAATTACAGTCGAATGGACAACGTCGTCG | CCGCTTTACTTGTACTCAGATCTTCTTGAGAA | |
| *CYP72A252v1* | *E. phyllopogon* | TACAATTACAGTCGAATGGCGGCTCGCCTTCT | CCGCTTTACTTGTACTCAGAGCTTCTTCAATCTGA | |
| *CYP72A252v2* | *E. phyllopogon* | TACAATTACAGTCGAATGGCGGCTCGCCTTCT | CCGCTTTACTTGTACTCACGATCCGAGCTTCTTCA | |
| *CYP78C* | *E. phyllopogon* | TACAATTACAGTCGAATGACCTTAATCCCGGC | CCGCTTTACTTGTACTCACACGCCGCGCCTC | |
| *CYP92A198* | *E. phyllopogon* | TACAATTACAGTCGAATGATGGAGTTCGCACCT | CCGCTTTACTTGTACTCACAGCCCAGCATAGA | |
| *CYP81A12* | *E. phyllopogon* | TACAATTACAGTCGAATGGATAAGGCCTACGTGGC | CCGCTTTACTTGTACGATGTTCTTCAGGAGCTCTGA | |
| **pYeDP60 cloning** | | | | |
| *CYP709C69* | *E. phyllopogon* | CCCGGGTACCAAAAAAATGGGTCTGGCTTGGA | ATCCCCCGCGGAATTCTTACACCTCCACGCGC | |
| *CYP709C5* | Rice | CCCGGGTACCAAAAAAATGGGTAATCTGGGGTGGAT | ATCCCCCGCGGAATTCCTACACCTTGAGACTCTTGAGGA | |
| *CYP709C9* | Rice | CCCGGGTACCAAAAAAATGGCCATGGGATTGTTAGC | ATCCCCCGCGGAATTCCTACAGCTTGAGGCTCTTGAGG | |
| **qPCR for *Arabidopsis thaliana*** | | | | |
| *CYP709C69*[b] | pCAMBIA1390 vector | ATCCTGTTGCCGGTCTTG | CCATCTCATAAATAACGTCATGC | |
| *GAPDH*[c] | *A. thaliana* | TTGGTGACAACAGGTCAAGCA | AAACTTGTCGCTCAATGCAATC | |
| **qPCR for *E. phyllopogon*** | | | | |
| *CYP71C35* | *E. phyllopogon* | CTGTCCAACCTCATCTACCAC | AAAATCTAGACGGTTAGGCCAAG | 200 nM |
| *CYP71E* | *E. phyllopogon* | CGAACACGCCAGATTGCTAC | CATAACTATTCGTCGGATCAAATTC | 300 nM |
| *CYP71S* | *E. phyllopogon* | CTGCCATGAAAGGGACGTAGT | GTAAAGGTACAAAACACGATAGACG | 300 nM |
| *CYP72A122*[d] | *E. phyllopogon* | CATGGCGCACCAATTATTC | TGCACTGTGACATACTCATTGAC | 200 nM |
| *CYP72A252v1* | *E. phyllopogon* | TATAGCAGCCTGTTCGGTTG | TCGTGTACACTGCTTCCTGTG | 200 nM |
| *CYP72A252v2* | *E. phyllopogon* | AGAGCGTGCATGAATAGGAAG | GTGATCAAAACAAGGGCTTAGAG | 200 nM |
| *CYP88A* | *E. phyllopogon* | TTCACAGATAGCATAGCACCAAC | GCCAAGATCACCAAAGTCTCC | 300 nM |
| *CYP709C67* | *E. phyllopogon* | GATTGGAAGCCATTCTTGTGA | CACACCGCGAAACAAAATTA | 300 nM |
| *CYP709C69* | *E. phyllopogon* | CGCGTGGAGGTGTAAGAATG | TAACATAAGCTACACGCATGACC | 300 nM |
| *CYP709C70* | *E. phyllopogon* | AAGCTTCTCTGTATTGTACGTAGTGC | GCGGTATATTAGTAATTCATTGAAAA | 300 nM |
| *EIF4B**** | *E. phyllopogon* | CCAGTCCCTTTTTGTTTTGGA | CTACAGCATACAGAGGTGATCAAT | 100 nM |

a, Primer concentration in real-time PCR. b, Developed in Tanigaki et al., (2021). c, Developed in Czechowski et al., (2005). d, Developed in Iwakami et al., (2014b).
