## Supplementary material for "Genetically-linked simultaneous overexpression of multiple herbicide-metabolizing genes for broad-spectrum resistance in an agricultural weed *Echinochloa phyllopogon*": Table S2

### Table S2. The summary of herbicides used for the functional characterizations of *CYP709C69*

| Group | Mode of Action | Chemical Group | Herbicide | Doses | | | |
|---|---|---|---|---|---|---|---|
| 1 | Inhibition of Acetyl CoA Carboxylase | Aryloxyphenoxy-propionate | Clodinatop-propargyl | 0 nM | 40 nM | 80 nM | 120 nM |
| | | | Cyhalofop-buthyl | 0 μM | 1 μM | 3 μM | 10 μM |
| | | | Fenoxaprop-P-ethyl | 0 μM | 100 nM | 300 nM | |
| | | | Quizalofop-P-ethyl | 0 μM | 2 μM | 4 μM | |
| | | | Metamifop | 0 μM | 0.3 μM | 1 μM | 3 μM |
| | | Cyclohexanediones | Tralkoxydim | 0 μM | 0.3 μM | 0.6 μM | |
| | | *Not annotated* | Pinoxaden | 0 nM | 50 nM | 100 nM | |
| 2 | Inhibition of Acetolactate Synthase | Pyrimidinyl benzoates | Pyriftalid | 0 nM | 1 nM | 10 nM | 30 nM |
| | | | Bispyribac-sodium | 0 nM | 10 nM | 30 nM | 100 nM |
| | | Sulfonanilides | Pyrimisulfan | 0 nM | 0.3 nM | 1 nM | 3 nM |
| | | Imidazolinone | Imazamox | 0 nM | 1 nM | 10 nM | 30 nM |
| | | Triazolinones | Propoxycarbazone-sodium | 0 nM | 0.1 nM | 1 nM | 10 nM |
| | | Triazolopyrimidine – type 2 | Penoxsulam | 0 nM | 0.1 nM | 1 nM | 3 nM |
| | | Sulfonilurea | Azimsulfuron | 0 nM | 0.1 nM | 0.3 nM | 1 nM |
| | | | Pyrazosulfuron-ethyl | 0 nM | 0.1 nM | 0.3 nM | 1 nM |
| | | | Chlorsulfuron | 0 nM | 1 nM | 3 nM | 10 nM |
| | | | Propyrisulfuron | 0 nM | 0.3 nM | 1 nM | 3 nM |
| | | | Bensulfuron-methyl | 0 nM | 0.1 nM | 1 nM | 3 nM |
| 3 | Inhibition of Microtubule Assembly | Dinitroanilline | Trifluralin | 0 μM | 1 μM | 3 μM | 10 μM |
| 4 | Auxin Mimics | Phenoxy-carboxylates | 2,4-D | 0 nM | 30 nM | 100 nM | 300 nM |
| | | Quinoline-carboxylates | Quinclorac | 0 μM | 10 μM | 60 μM | 300 μM |
| 5 | D1 Serine 264 binders (and other non-histidine 215 binders) | Amide | Propanil | 0 μM | 1 μM | 10 μM | 100 μM |
| | | Triazines | Cyanazine | 0 μM | 0.3 μM | 1 μM | 3 μM |
| 6 | D1 Histidine 215 binders | *Not annotated* | Bentazone | 0 μM | 1 μM | 3 μM | 30 μM |
| | | Nitriles | Bromoxynil | 0 μM | 1 μM | 3 μM | 30 μM |
| 9 | Inhibition of Enolpyruvyl Shikimate Phosphate Synthase | *Not annotated* | Glyphosate | 0 μM | 1 μM | 10 μM | 100 μM |
| 10 | Glutamine Synthetase | Phosphinic acids | Glufosinate | 0 μM | 10 μM | 30 μM | 100 μM |
| 12 | Inhibition of Phytoene Desaturase | Pheyl-ethers | Diffufenican | 0 nM | 1 nM | 3 nM | 10 nM |
| | | N-Phenyl heterocycles | Norflurazon | 0 nM | 30 nM | 100 nM | 300 nM |
| | | Diphenyl heterocycles | Fluridone | 0 nM | 3 nM | 10 nM | 30 nM |
| 13 | Inhibition of Deoxy-D-Xyulose Phosphate Synthase | Isoxazolidinones | Clomazone | 0 nM | 100 nM | 300 nM | 1000 nM |
| 14 | Inhibition of Protoporphyrinogen Oxidase | *Not annotated* | Pyracionil | 0 nM | 0.3 nM | 1 nM | 3 nM |
| | | N-Phenyl-imides | Flumioxazin | 0 nM | 0.1 nM | 1 nM | 10 nM |
| | | N-Phenyl-oxadiazolones | Oxadiargyl | 0 nM | 100 nM | 300 nM | 600 nM |
| 15 | Inhibition of Very Long-Chain Fatty Acid Synthesis | α-Chloroacetamides | Butachlor | 0 μM | 3 μM | 6 μM | 20 μM |
| | | | Pretilachlor | 0 μM | 3 μM | 10 μM | 30 μM |
| | | Thiocarbamate | Molinate | 0 nM | 60 nM | 100 nM | 300 nM |
| | | | Thiobencarb | 0 μM | 10 μM | 30 μM | 100 μM |
| | | Isoxazolines | Pyroxasulfone | 0 μM | 1 μM | 10 μM | 30 μM |
| | | | Fenoxasulfone | 0 μM | 1 μM | 100 μM | 300 μM |
| 22 | PS I Electron Diversion | Pyridiniums | Paraquat | 0 nM | 30 nM | 100 nM | 300 nM |
| 27 | Inhibition of Hydroxyphenyl Pyruvate Dioxygenase | Triketone | Mesotrione | 0 nM | 0.1 nM | 1 nM | 10 nM |
| 29 | Cellulose Synthesis | Benzamide | Isoxaben | 0 nM | 1 nM | 10 nM | 100 nM |
| | | Nitriles | Dichlobenil | 0 μM | 0.3 μM | 1 μM | 3 μM |
| Not classified | Unknown | *Not annotated* | Pyributicarb | 0 nM | 100 nM | 300 nM | 1,000 nM |
| | | *Not annotated* | Bromobutide | 0 μM | 3 μM | 10 μM | 30 μM |

The herbicide classifications follow the HRAC system (https://www.hracglobal.com/). Another mode of action was annotated for quinclorac (cellulose synthesis) and bromoxynil (uncouplers).
